# Non-canonical function of ADAM10 in presynaptic plasticity

**DOI:** 10.1101/2024.05.12.593731

**Authors:** Julia Bär, Tomas Fanutza, Christopher C. Reimann, Lisa Seipold, Maja Grohe, Janike Rabea Bolter, Flemming Delfs, Michael Bucher, Christine E Gee, Michaela Schweizer, Paul Saftig, Marina Mikhaylova

**Affiliations:** AG Optobiology, Institute of Biology, Humboldt Universität zu Berlin, Berlin, 10115 Germany; Guest Group “Neuronal Protein Transport”, Center for Molecular Neurobiology, University Medical Center Hamburg-Eppendorf, Hamburg, 20251 Germany; discontinued in August 2023; Biochemisches Institut, Christian Albrechts-Universität Kiel, Kiel, 24098 Germany; Department of Synaptic Physiology, Center for Molecular Neurobiology, ZMNH, University Medical Center Hamburg-Eppendorf, 20251 Hamburg, Germany; Morphology and Electron Microscopy, University Medical Center Hamburg-Eppendorf, Center for Molecular Neurobiology, ZMNH, Hamburg, 20251 Germany

**Keywords:** ADAM10, mossy fiber facilitation, protease, vesicle release

## Abstract

A Disintegrin And Metalloproteinase 10 (ADAM10) plays a pivotal role in shaping neuronal networks by orchestrating the activity of numerous membrane proteins through the shedding of their extracellular domains. Despite its significance in the brain, the specific cellular localization of ADAM10 remains not well understood due to a lack of appropriate tools. Here, using a specific ADAM10 antibody suitable for immunostainings, we discover that ADAM10 is localized to presynapses and especially enriched at presynaptic vesicles of mossy fiber (MF)-CA3 synapses in the hippocampus. These synapses undergo pronounced frequency facilitation of neurotransmitter release, a process that play critical roles in information transfer and neural computation. We demonstrate, that in conditional ADAM10 knockout mice the ability of MF synapses to undergo this type of synaptic plasticity is greatly reduced. The loss of facilitation depends on the cytosolic domain of ADAM10 and association with the calcium sensor synaptotagmin 7 rather than its proteolytic activity. Our findings unveil a new pathway contributing to the regulation of synaptic vesicle exocytosis.

## Introduction

A Disintegrin And Metalloprotease 10 (ADAM10) is prominently expressed in the brain [24]. ADAM10 has received much attention due to its role in Alzheimer’s (AD), prion disease and Huntington’s disease (HD) [2, 27, 55]. It catalyzes ectodomain shedding, i.e. the proteolytic release of an ectodomain, from several important cell surface proteins such as amyloid precursor protein (APP), Notch receptor and prion protein [40]. ADAM10 itself is a membrane spanning protein with a short cytosolic domain, a transmembrane region and an ectodomain that harbors the catalytic site. Although much is known about ADAM10 substrates and its involvement in pathological processes, the physiological functions of ADAM10 in the brain are not fully understood. One of the limiting factors is the absence of suitable tools to label endogenous ADAM10, along with the tightly regulated processing and trafficking of ADAM10. These factors constrain the feasibility of overexpression and mutagenesis studies. Due to defective Notch signaling, ADAM10-deficient mice die before birth [13]. However, depleting ADAM10 postnatally by crossing floxed ADAM10 mice with nestin or CaMKIIα-Cre driver lines allowed generation of neuron-specific conditional ADAM10 knockouts (ADAM10 cKO). These lines are used to study the role of ADAM10 in the homeostasis of adult neuronal networks [21, 36]. Mechanistically, ADAM10’s functions are mainly linked to the shedding of unique postsynaptic substrates, in particular synaptic cell adhesion molecules such as neuroligin-1, N-cadherin, NCAM, Ephrin A2 and A5 [26] which are of importance for the establishment and maintenance of dendritic spines in the CA1 hippocampal region [36, 46]. Taken together, there is an imminent need for tools that would facilitate the investigation of ADAM10 functions in the brain.

In this study we employed a new anti-ADAM10 antibody to systematically investigate the localization of ADAM10 and its functions within specific cell compartments. Using electron- and super-resolution microscopy we found that ADAM10 is enriched in vesicles of excitatory and inhibitory presynaptic sites. Especially prominent expression was seen in hippocampal mossy fiber (MF)-CA3 terminals. Short-term plasticity of MF-CA3 synapses is pivotal for information processing and network computation [12]. We reasoned that these synapses could serve as a suitable model for studying the physiological functions of ADAM10. We discovered that ADAM10 depletion strongly reduces facilitation at MF-CA3 synapses without gross changes in synaptic topology or ultrastructure. In contrast to known roles of ADAM10 [26, 40], the impairment of presynaptic facilitation is independent of proteolytic activity. Instead, the cytosolic domain of ADAM10 and an association with synaptotagmin 7 (syt7) is required. Syt7 is a calcium sensor essential for presynaptic facilitation of vesicle release [20]. Knockout of syt7 phenocopies the MF-CA3 phenotype observed in slices from ADAM10 cKO mice. Interference with syt7 and ADAM10 association using a tat-peptide reduces presynaptic facilitation at MF-CA3 synapses in slices of wildtype mice. Thus, ADAM10 appears to tune short-term synaptic plasticity in a non-proteolytic way by affecting syt7 function in presynaptic terminals.

## Materials and methods

### Reagents and resources

Information regarding essential tools is provided in the table below.

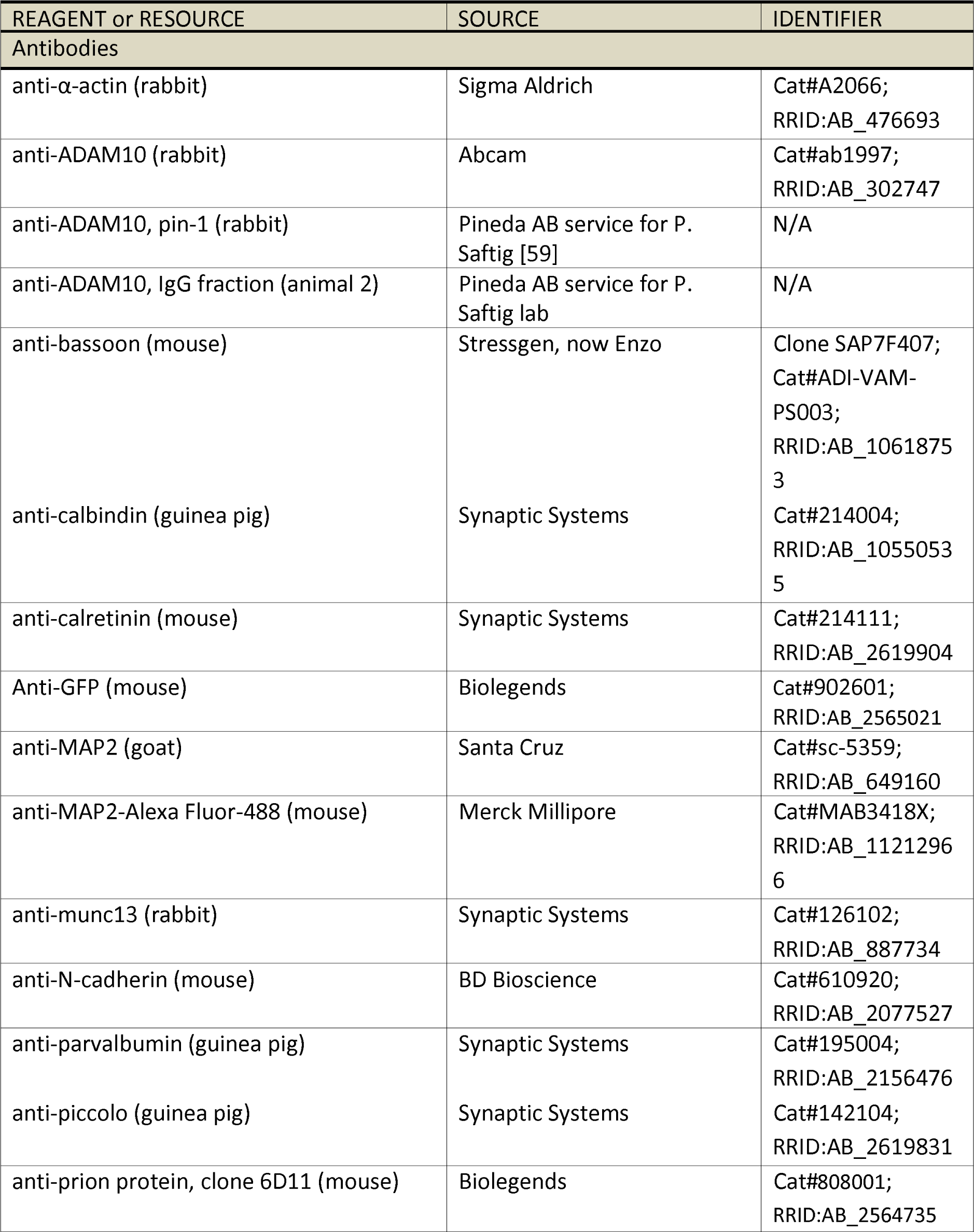
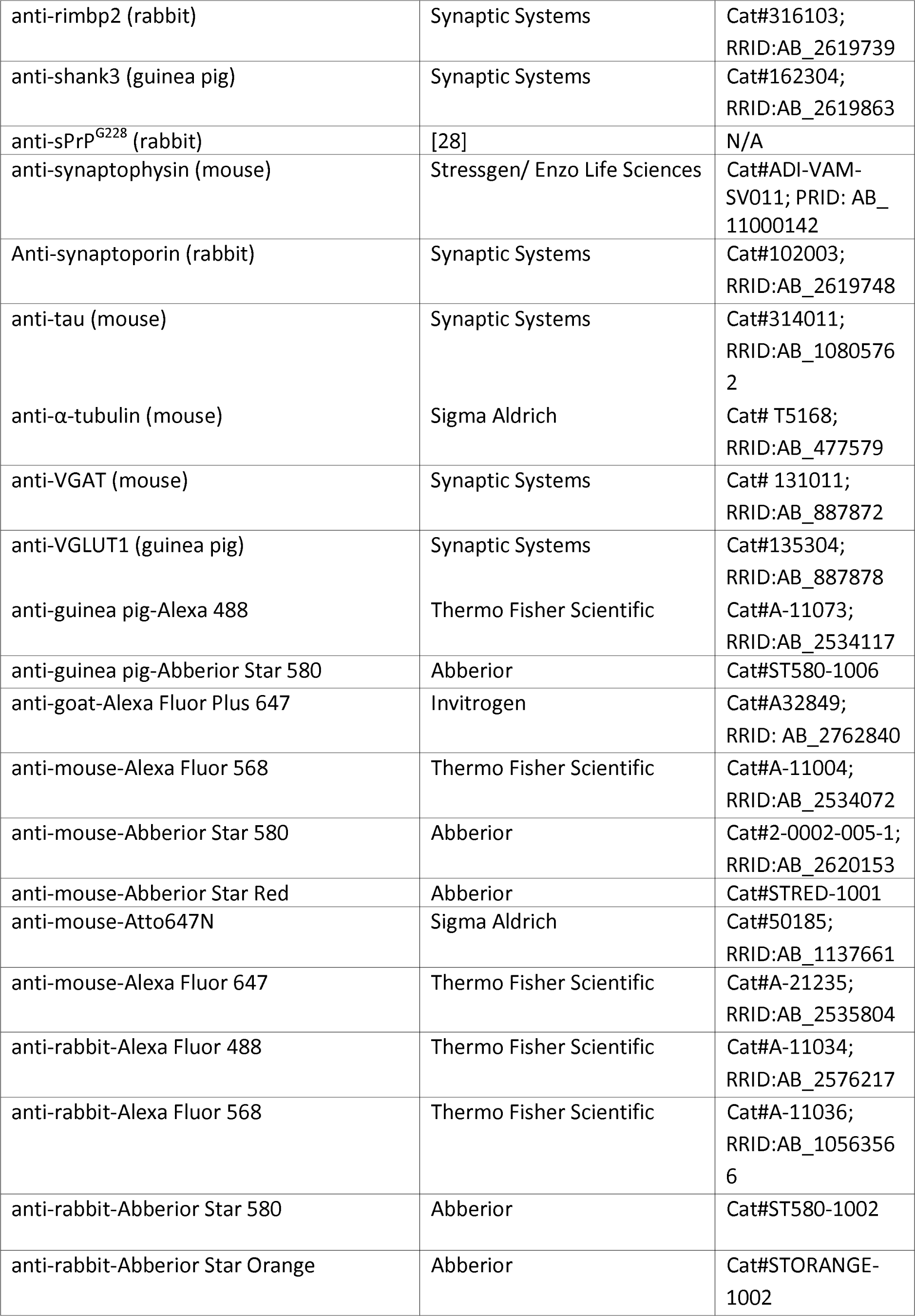
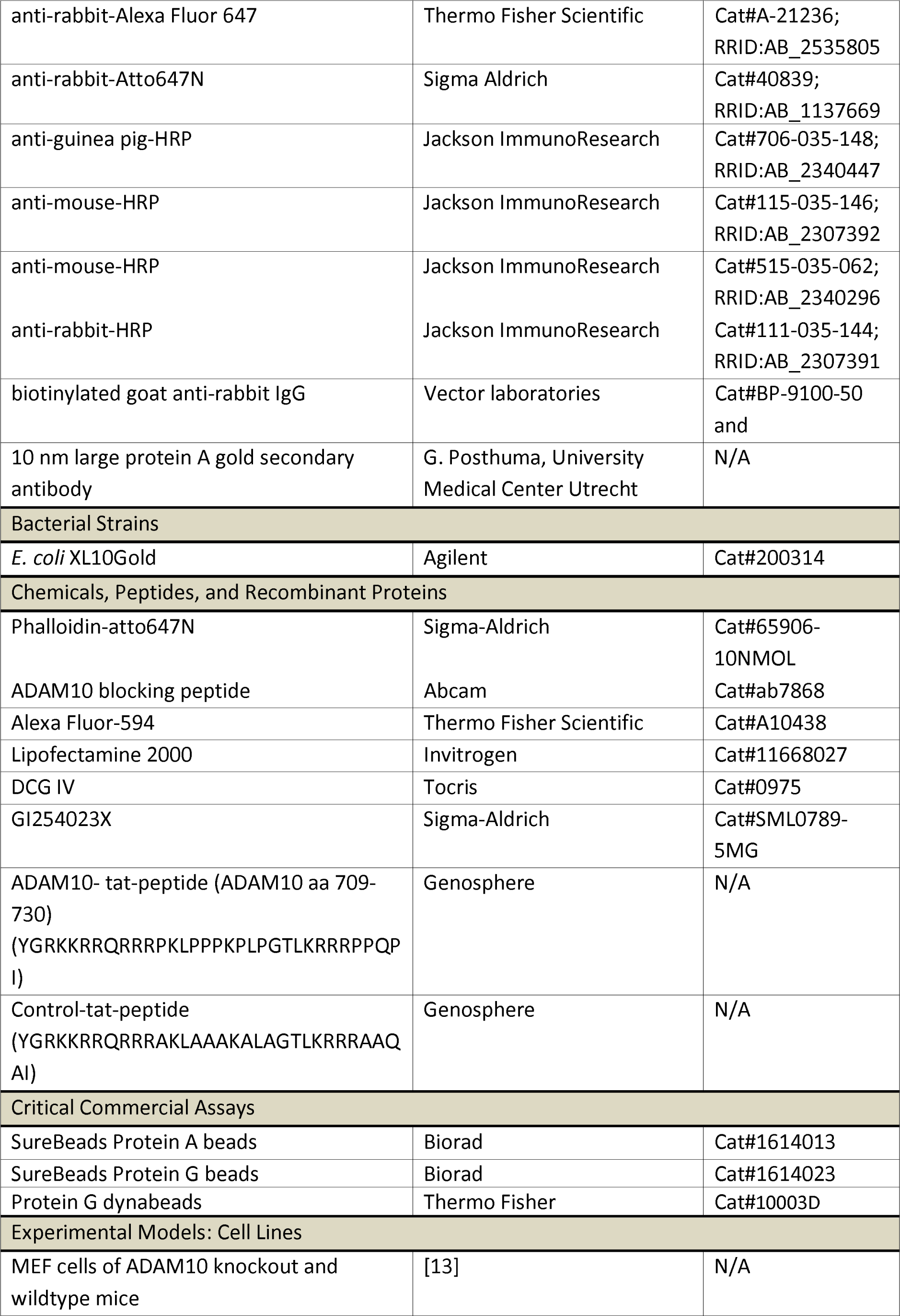
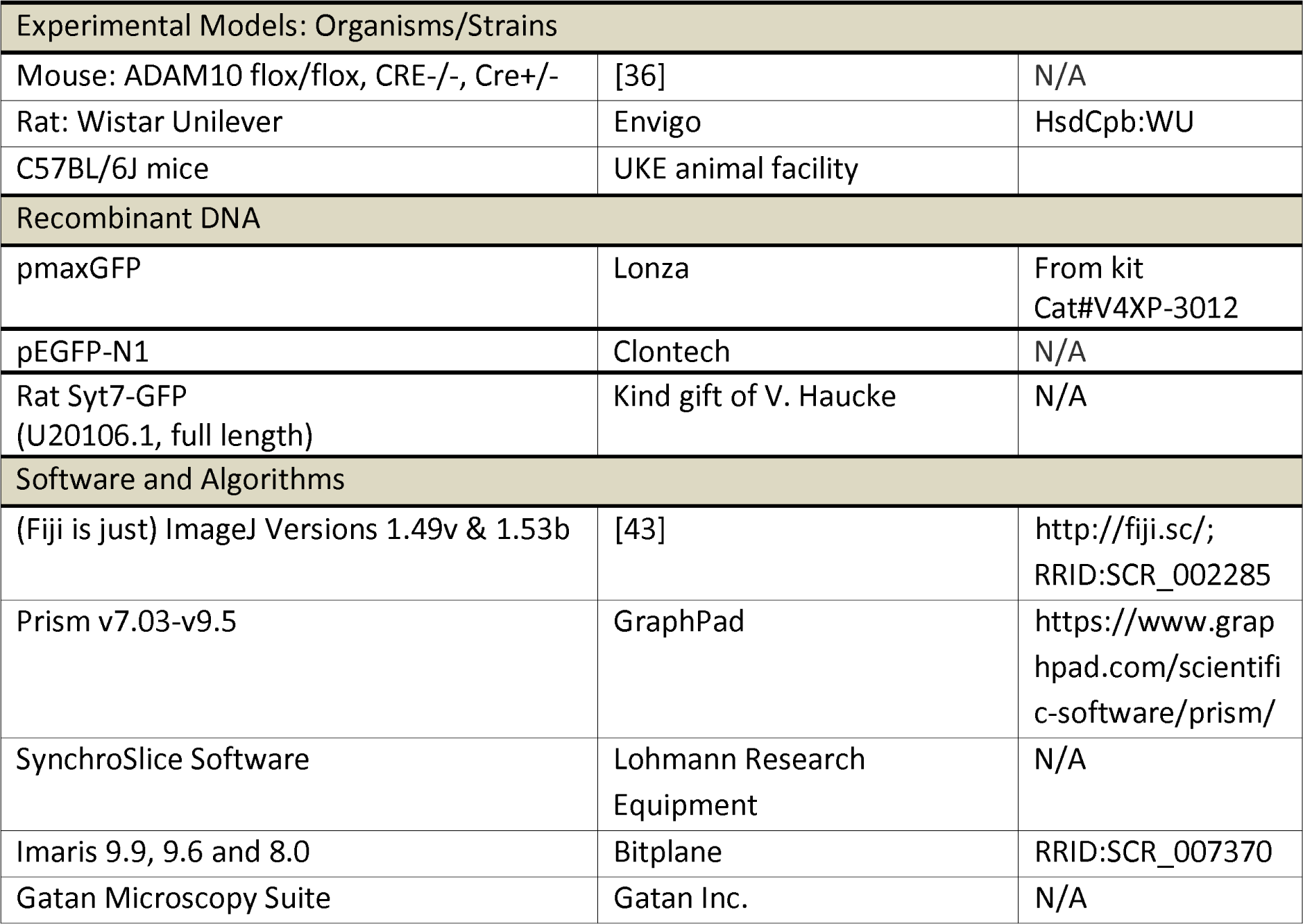

### Animals

Conditional ADAM10 knockout mice (cKO; ADAM10^flox/flox^ Cre^+/-^) and littermate controls (ADAM10^flox/flox^ Cre^-/-^) [36] were obtained by breeding ADAM10^flox/flox^ Cre^-/-^ or ADAM10^flox/+^ Cre^-/-^ with ADAM10^flox/+^ Cre^+/-^ mice. Male and female P21-23 mice were used for this study.

Mice were bred and maintained at CAU, Kiel, Schleswig-Holstein. Animals were housed in individually ventilated cages with a capacity of five animals per cage. The room temperature was kept at 19-22 °C and humidity at 45-60 %. Alternating light-dark cycles of 12 h were applied and animals were fed with water and laboratory animal food (Ssniff Spezialdiäten) *ad libitum*.

Rats were obtained from the animal facility of the University Medical Center Hamburg-Eppendorf, UKE, Hamburg, Germany, approved by the local authorities of the city-state Hamburg (Behörde für Gesundheit und Verbraucherschutz, Fachbereich Veterinärwesen) and the animal care committee of the University Medical Center Hamburg-Eppendorf.

### Peptides

Custom tat-peptides targeting amino acids 709-730 of ADAM10 C-terminus (NP_031425.2) and control peptide (prolines changed to alanines) with >95 % purity (HPLC, mass spectrometry and solubility tested) were from Genosphere Biotechnologies (France).

### Biochemical methods

#### Cell lysates, tissue extracts, and immunoblotting

Freshly dissected cortices of P21 ADAM10 cKO and littermate controls were shortly rinsed in PBS, shock frozen in liquid nitrogen and stored until use at -80 °C. Tissue was manually homogenized using 26G and 20G needles in RIPA buffer [50 mM Tris/HCl pH 8, 150 mM NaCl, 0.1 % sodium dodecyl sulfate (SDS), 0.5 % Triton-X-100, 0.5 % sodium deoxycholate and complete protease inhibitor cocktail (Roche)] at a 10 ml/g ratio. After incubation on ice for 20 min and centrifugation for 20 min at 20000 x g supernatant was collected. After addition of 4x SDS sample buffer (250 mM Tris pH 8.5, 8 % SDS, 40 % glycerol, 2 mM ethylenediaminetetraacetic acid (EDTA), 0.01 % bromophenol blue, 100 mM dithiothreitol) further dilution with 1x SDS sample buffer and boiling for 5 min, samples were separated on a 10 % polyacrylamide gel, and transferred to a nitrocellulose membrane using a wet blot system.

Cell homogenates of ADAM10 wt and KO mouse embryonic fibroblasts were prepared as follows. Cells were shortly washed with warm TBS, harvested in hot 1x SDS sample buffer and boiled for 5 min. Equal amount of sample were separated on a 4-20 % polyacrylamide gel and transferred to a PVDF membrane. All membranes were blocked for 1 h in 5 % milk in Tris-buffered saline (TBS, 20 mM Tris, 150 mM NaCl, pH 7.4,) with 0.1 % Tween-20 (TBST) and incubated overnight at 4 °C with primary antibodies in TBS with 0.02 % NaN_3_. After washing in TBST and TBS corresponding membranes were incubated with secondary antibodies in 5 % milk in TBST for 1h, washed, and signals detected using ECL solution on an INTAS ChemoCam Imager (Intas Science Imaging). α-tubulin was used as loading control. Following electrophysiological recordings (see below) slices of ADAM10 wt and cKO mouse brains and wildtype slices from the ADAM10 inhibitor experiment were homogenized with 400 µl RIPA lysis puffer (20 mM Tris/HCl pH 7.4, 150 mM NaCl, 1 mM EDTA, 1 mM EGTA, 2.5 mM Sodium deoxycholate, 1 % (v/v) NP-40 and protease inhibitor cocktail (Roche)) and two ceramic beads using the Precellys tissue homogenizer (Bretin Technologies) with two cycles of 30 s at 6500 rpm and a 30 s pause. Homogenates were kept on ice for 1 h and centrifuged for 10 min at 13000 x g and 4 °C. Supernatant was collected, diluted with 5x SDS loading buffer (625 mM Tris/HCl pH 6.8, 10 % SDS, 50 % glycerol, 200 mM dithiothreitol, 0.01 % bromophenol blue) and incubated at 60 °C for 20 min. For immunoblotting samples were separated on a 10 % SDS polyacrylamide gel and transferred to a nitrocellulose membrane using the wet blot system with a constant amperage of 450 mA (for N-cadherin) or 800 mA (for other proteins) for 2 h. Membranes were blocked for 1 h in 5 % bovine serum albumin (BSA,for N-cadherin) or 5 % milk powder in TBST and incubated with primary antibodies prepared in 5 % milk powder in TBST overnight at 4 °C. Membranes were washed in TBST and incubated with secondary antibodies diluted in 5 % milk powder in TBST for 1 h at room temperature. Detection of signals was performed using ECL solution on an Amersham Imager A 680 (GE Healthcare). Information regarding antibodies and other reagents is provided in Reagents and resources table.

#### Synaptosome preparation

Synaptosomes were prepared from 4 male C57BL6J mice aged 12-14 weeks as described previously [47] with slight modifications: All steps were performed at 4°C or on ice. Forebrain and midbrain (without olfactory bulb) were dissected, rinsed in buffer A (0.32 M sucrose, 5 mM HEPES, protease inhibitors, pH 7.4) and homogenized in 10 mg/ml buffer A by 12 strokes at 900 rpm with a motor-operated homogenizer. Homogenates were centrifuged at 1000 x g for 10 min to obtain pellet 1 and supernatant 1. Pellet 1 was again solubilized in buffer A, homogenized and centrifuged. The resulting supernatant was combined with supernatant 1 and used for further fractionation. The combined supernatants were centrifuged at 12000 x g for 20 min to obtain pellet 2, which was again rehomogenized in buffer A by 6 strokes at 900 rpm. Centrifugation at 12000 x g for 20 min obtained the crude membrane fraction (pellet P2‘). Pellet P2’ was solubilized in 1.5 ml/g buffer B (0.32 M sucrose, 5 mM Tris, protease inhibitors, pH 8.1) loaded on a 0.85 M, 1 M, 1.2 M sucrose (in 5 mM Tris pH 8.1) gradient, centrifuged at 85000 x g for 2 h. The synaptosome fraction was collected at the interface of the 1 and 1.2 M sucrose phase, immediately shock frozen in liquid nitrogen and stored at -80 °C until further use.

#### Endogenous co-immunoprecipitations

For immunoprecipitation of ADAM10, synaptosome preparations were centrifuged for 30 min at 100.000 x g. The supernatant was discarded and synaptosomes lysed in EBC buffer (120 mM NaCl, 50 mM Tris-HCl, pH 7.4, 1 % NP-40, protease inhibitor cocktail) by two times sonification for 15 s followed by 30 min incubation on ice and another two times sonification for 15 s. After centrifugation for 10 min at 14000 x g, supernatants were collected, and total protein concentration was determined. 50 µl of the homogenates were kept as input control and incubated with SDS-loading buffer for 20 min at 60 °C. Remaining homogenates were adjusted to equal total protein amounts and split into calcium and calcium-free conditions by addition of 200 µM CaCl_2_ or 2 mM EGTA. These conditions were kept throughout the experiment in the EBC lysis buffer. Preclearing of homogenates was performed. Per sample 50 µl of Protein G Dynabeads^TM^ (Invitrogen) were two times washed with EBC lysis buffer, added to the homogenates and incubated for 30 min at 4 °C on a rotating wheel. Supernatants were collected and 2 µl of the anti-ADAM10 Pin 1 antibody were added. EBC buffer only supplemented with anti-ADAM10 Pin 1 antibody served as antibody control, homogenate without ADAM10 antibody was used to control for unspecific protein binding to the beads (bead control). All samples were incubated overnight at 4 °C under constant rotation. At the same time Protein G Dynabeads^TM^ were prepared for immunoprecipitation. Per sample 80 µl of Protein G Dynabeads^TM^ were 2 times washed with EBC lysis buffer and blocked overnight at 4 °C under constant rotation with SEA BLOCK blocking buffer (Thermo Fisher). After overnight incubation, blocked Protein G Dynabeads^TM^ were two times washed and resuspended with EBC buffer. Immunoprecipitation samples were incubated with Protein G Dynabeads^TM^ for 30 min at room temperature under constant rotation. After five times washing with 500 µl EBC buffer for 5 min under constant rotation, supernatants were discarded. For immunoblot analysis beads were incubated with 1x SDS loading buffer for 20 min at 60 °C for and supernatants were collected. Information regarding antibodies and other reagents is provided in Reagents and resources table.

#### Heterologous CoIP from Neuro-2a cells

Neuro-2a cells were transfected with GFP or syt7-GFP for 24-48h using MaxPEI. Cells were harvested in culture medium, spun down, pellets washed in washing buffer (120 mM NaCl, 50 mM Tris-HCl, pH 7.4) and subsequently lysed in EBC lysis buffer containing either 200 µM CaCl_2_ or 2 mM EGTA using small needles and rest for 1 h on ice. After centrifugation at 20000 x g for 10 min at 4 °C supernatants were collected. CoIP was performed essential as described above (endogenous CoIP), omitting the preclearing step, using 1 µl antibody per IP and 25 µl of a 1:1 mix of protein A and G beads and with addition of either 200 µM CaCl_2_ or 2 mM EGTA to the EBC lysis buffer in all steps. Information regarding antibodies and other reagents is provided in Reagents and resources table.

#### Cell Culture

ADAM10 WT and KO mouse embryonic fibroblast (MEF) cells [21] and N2a cells were cultured in full medium (DMEM including 2 mM glutamine, supplemented with 10 % FCS and 100 U/ml penicillin/streptomycin). For immunostainings, MEFs were plated on gelatin-coated glass coverslips in full medium. Primary rat hippocampal neurons were prepared and maintained as described previously [51]. At div17, neurons were transfected with pmaxGFP using lipofectamine 2000 according to manufacturer’s instructions and fixed 24 hours later with 4% PFA/ 4% sucrose in PBS for 10 min.

#### Immunocytochemistry (ICC)

ICC on MEF cells and primary neurons was performed as described previously [5]. For the experiment with the ADAM10 antibody blocking peptide, a preincubation of antibody and blocking peptide at a 1:1 molar ratio for 1 h at 4 °C in PBS was performed before applying it to the cells. Information regarding antibodies and other reagents is provided in Reagents and resources table.

#### Perfusion

P19-21 cKO and wt mice were anesthetized by intra-peritoneal injection of 10 µl per g body weight of Ketamin (10 mg/ml) and Rompun (6 mg/ml) in a 0.9 % NaCl solution, transcardially perfused with 15 ml 0.1 M phosphate buffer (PB), fixed with 15 ml 4 % PFA + 1 % glutaraldehyde (GA) in 0.1M PB (for immunogold electron microscopy (EM)), 4 % PFA + 0.1% GA in 0.1M PB (for serial block face EM) or 4 % PFA (for fluorescent immunohistochemistry). Postfixation was performed over night at 4°C in 2% PFA (for Immunogold EM), 4 % PFA + 1% GA (for SBEM) or 4% PFA (for fluorescent IHC) in PB. Adult (7.5 month) wt mouse was anesthetized and perfused transcardially with 4 % (w/v) PFA in 0.1 M PB. The brain was dissected and postfixed for 4 h with 1 % glutaraldehyde/4 % (w/v) PFA in 0.1 M PB.

#### Cryo-sections and immunohistochemistry (IHC)

Perfused brains prepared for IHC were placed through 0.5 M and 1M sucrose solution in 0.1 M PB, frozen in -50 °C 2-methylbutane and stored at -80 °C until use. Sagittal cryosections of 40 µm thickness were prepared on a Zeiss Hyrax C 60 cryostat and sections stored in PBS. Immunohistochemistry was mainly performed as in [31]. In general, all incubation and washing steps are performed on a shaker to allow proper blocking, washing and antibody distribution. All antibody dilutions were prepared in IHC-blocking buffer (BB, PBS including 0.3 % Triton X-100, 10 % normal goat serum). Brain sections were incubated in IHC-BB for 30 min at RT. Incubation with primary antibody was performed at 4 °C for approximately 36 h-48 h. Afterwards, sections were first washed in PBS for 1h and then thoroughly washed in 0.2 % BSA in PBS at RT for at least 30 min with several changes of washing buffer. Incubation with fluorescently labelled secondary antibodies was done for 2h at RT. After final 3 x 10 min washing in PBS, and 10 min incubation with 1µg/ml DAPI in PBS, sections were rinsed in tap water, placed on superfrost objective slides (Superfrost ultra plus, Thermo Scientific), air dried and mounted with mowiol. Information regarding antibodies and other reagents is provided in Reagents and resources table.

#### Electrophysiological Recordings

ADAM10 cKO and wt littermates, and C57BL6/J (for GI254023X and tat-peptide experiments) male mice aged 3 weeks were used. The experimenter was blind to the genotypes and identity of tat-peptide. Animals were anesthetized with CO_2_ and decapitated. Brains were rapidly removed from the skull and placed in an ice-cold modified artificial cerebrospinal fluid solution (aCSF) containing (in mM): 110 choline chloride, 25 NaHCO_3_, 25 D-glucose, 11.6 Na-L-ascorbate, 7 MgSO_4_, 1.25 NaH_2_PO_4_, 2.5 KCl, 0.5 CaCl_2_ (pH 7.4 equilibrated with 95 % O_2_ and 5 % CO_2_). 400 µm thick transversal brain slices were prepared with a microtome vf-200 tissue slicer (Compresstome, Precisionary Instruments, USA) and then incubated at 30 °C for about 90 minutes in a physiologic aCSF containing (in mM): 124 NaCl, 26 NaHCO_3_, 10 D-glucose, 1 MgSO_4_, 1 NaH_2_PO_4_, 4 KCl, 2.4 CaCl_2_ (pH 7.4 equilibrated with 95 % O_2_ and 5 % CO_2_). After the incubation time, the hemi-slices were transferred to Synchroslice recording chambers (Lohmann Research Equipment) perfused with aCSF at a flow rate of ∼2 ml/min using a peristaltic pump (minipulse 3, Gilson, USA). The experiments were performed at 30 °C. For the experiments with ADAM10 inhibitor GI254023X, 20 µM inhibitor was included during the incubation time for up to 180 minutes. For tat-peptide experiments, 3 µM peptides were included during incubation time and recordings. All recordings were executed by using a 2-channel Miniature Preamplifier (multichannel systems, Germany). The field excitatory post synaptic potentials (fEPSPs) extracellular recordings were performed by using a single fiber electrode (Lohmann Research Equipment, Germany) placed in the CA3 pyramidal cell body layer. fEPSPs were evoked by stimulating the mossy fibers using a semi-micro concentric bipolar electrode (Lohmann Research Equipment, Germany) placed nearby the granule cell layer or in the hilus region. Square-wave current pulses were generated by a stimulus generator (Multichannel Systems STG 4008, Germany) and delivered to the tissue. The recorded MF-CA3 signal was routinely verified by the following procedures, and the experiments were discarded if they did not match the criteria. 1) Synaptic response amplitudes were largely increased when the frequency facilitation protocol was applied (5 pulses, frequency 20 Hz). 2) fEPSPs were only accepted if responses showed an exponential increase in synaptic facilitation after each pulse. 3) Application of the group II mGluR agonist (2S,211R)-2-(211,311-Dicarboxycyclopropyl) glycine (DCG-IV) (1 μM) at the end of the experiment had to block synaptic transmission responses. Paired-pulse ratio (PPR) was measured by delivering two stimuli for five times 30 seconds apart at 50, 100, 200, 500 ms inter-stimulus intervals. The PPR was calculated by dividing the amplitude of the second EPSP by the amplitude of the first EPSP. Synaptic facilitation was examined by repetitive stimulation (5 times) for each inter-stimulus interval, and the resulting potentials were averaged. Frequency facilitation was measured by delivering five pulses for five times 30 seconds apart at 50 and 100 ms inter-stimulus intervals and potentials were averaged. The facilitation was calculated by dividing the amplitude of each EPSP response by the amplitude of the first EPSP response. All signals were low pass filtered at 211kHz and digitized at 1011kHz. Recordings were analyzed by using the SynchroSlice (Lohmann Research Equipment) software.

### Microscopy

#### Widefield microscopy

Widefield imaging was performed at a Nikon Eclipse Ti-E microscope controlled by VisiView software, with a 100x, 60x objective or 10x, yielding pixel sizes of 65 nm, 108 nm or 650 nm equipped with standard GFP, RFP, and Cy5 filters. Illumination was achieved by a LED light source. Images were taken at 16-bit depth and 2048 x 2048 pixel.

#### Spinning disc confocal microscopy

Spinning-disc confocal microscopy was performed with a Nikon Eclipse Ti-E controlled by VisiView software. Illumination was done by 488 nm, 561 nm, and 639 nm excitation lasers, coupled to a CSU-X1 spinning disk unit via a single-mode fiber. Emission was collected through a quad band filter (*Chroma*, ZET 405/488/561/647m) on an Orca flash 4.0LT CMOS camera (*Hamamatsu*). Use of 100x TIRF objective (*Nikon*, ApoTIRF 100x/1.49 oil) achieved a pixel size of 65 nm, z step size was set to 0.3 µm.

#### Laser scanning confocal microscopy and STED

Fixed primary cultures were imaged at a Leica TCS SP5 confocal microscope (*Leica microsystems, Manheim, Germany*), controlled by Leica Application Suite Advanced Fluorescence software. Samples were imaged using a 63x oil objective (*Leica,* 63x HCX PL APO Lbd. Bl. Oil/1.40 oil). Fluorophores were excited with multi-Argon 488 nm, Diode Pumped Solid State 561 nm, HeNe 633 nm lasers and signals detected using HyD detectors. Pixel depth of 8-bit and frame averaging of 2 was used. For synapse imaging, dimensions were set to 512 x 512 pixels, a pixel size of 80 nm, and a z-step size of 0.29 was set for z-stacks. For neuronal overview images, dimension were 1024 x 1024 pixels, pixel size 240 nm and z-steps of 0.4 µm were set.

A Leica TCS SP8-3X gatedSTED system equipped with a 470 to 670 nm pulsed white light laser (WLL) and a 100x objective (*Leica,* HC APO CS2 100x/1.40 oil) was used for confocal and gated STED imaging. For excitation of the respective channels the WLL was set to 650 nm for Atto647N, 561 nm for Cy3, 580 nm for Abberior Star 580, and 488 nm for Alexa Fluor. Signals were detected using HyD detectors. Confocal images were acquired as single planes, with 18 nm pixel size, 8-bit image depth, and a line average 8. STED was obtained with a 775 nm pulsed depletion laser for Abberior Star 580 and Atto647N, and with a 592 nm continuous wave laser for Alexa Fluor 488. Emission spectra were collected between 660-730 nm, 580-620 nm, 500-530 nm. Detector time gates were set to 0.5-6 ns for Abberior Star 580/Atto647N and 1.5 ns-6 ns for Alexa Fluor 488. Images were acquired as single planes with a pixel size of 22.73 nm, 12-bit pixel depth, 600 lines per second and 16 x line averaging. The corresponding confocal were acquired with identical settings, except that detection time gates were set to 0.3 ns - 6 ns for all channels and the excitation power was reduced.

STED imaging of mossy fiber boutons (MFBs) in cryosections of ADAM10 wt and cKO mice was performed on an Olympus IX83 microscope equipped with STEDycon unit (Abberior) and pulsed diode laser 561 nm (for excitation of Abberior Star orange) and 640 nm (for excitation of Abberior Star Red) as well as 775 nm pulsed STED laser. Signals were detected with single photon counting avalanche photodiodes with single bandpass filters for 650-700 nm and 575-625 nm, respectively. Excitation of DAPI for orientation within the section was performed with a CW diode laser 405 nm, and detection of signals was achieved with a single photon counting avalanche photodiode with single bandpass filter for 420-480 nm. An UPLX APO 100x objective (1.45 NA) was used. Dual color images were taken as single planes with a pixel size of 25 nm, 16-bit pixel depth, 10 µs pixel dwell time, 10 x line accumulation. Large fields of view (up to 20 µm x 20 µm) were acquired.

### Electron microscopy

#### DAB and Immunogold

Perfused brains were dissected and 100 µm thick sagittal sections were cut with a Vibratome (Leica VT 1000S). Thereafter pre-embedding immunoelectron microscopy was performed as following: Vibratome sections with the hippocampal formation were selected and cryoprotected in 2.3 M sucrose and subjected to two cycles of freeze-thaw in liquid nitrogen. After rinsing in PBS, the sections were incubated with 10 % horse serum containing 0.2 % BSA (blocker) for 15 min and incubated with anti-ADAM10 antibody diluted 1:250 in PBS containing 1 % horse serum and 0.2 % BSA (carrier) over night. The sections were washed with PBS, then further incubated with biotinylated goat anti rabbit IgG (Vector laboratories, Burlingame, CA) diluted 1:1000 in carrier for 90 min. After rinsing, they were incubated with ABC (Vector Labs) diluted 1:100 in PBS for 90 min. Sections were washed again in PBS and reacted in diaminobenzidine (DAB)-H_2_0_2_ solution (Sigma St. Louis, USA) for 10 min. Thereafter the sections were rinsed three times in 0.1 M sodium cacodylate buffer (pH 7.2-7.4) (Sigma-Aldrich, Buchs, Switzerland) and incubated with 1 % osmium tetroxide (Science Services, Munich, Germany) in cacodylate buffer for 20 minutes on ice. The osmication of sections was followed by dehydration through ascending ethyl alcohol concentration steps and rinsed twice in propylene oxide (Sigma-Aldrich, Buchs, Switzerland). Infiltration of the embedding medium was performed by immersing the sections first in a mixture of 2:1 of propylene oxide and Epon (Carl Roth, Karlsruhe, Germany) then in a 1:1 mixture and finally in neat Epon and hardened at 60 °C for 48 h. For post-embedding immunogold labeling, small pieces of cryoprotected hippocampal CA3 region (2.3 M sucrose) were mounted on specimen holders immersed in liquid nitrogen and ultrathin sections (70 nm) were cut and labeled according to [44]. Briefly, sections were collected on Carbon-Formvar-coated nickel grids (Science Services GmbH, Germany). Rabbit anti ADAM10 antibody dilution 1:100 was recognized with 10 nm large protein A gold secondary antibody (purchased from G. Posthuma, University Medical Center Utrecht). Ultrathin sections were examined in an EM902 (Zeiss, Germany) or JEM-2100Plus (JEOL, Germany). Pictures were taken with a TRS 2K digital camera (A. Tröndle, Moorenweis, Germany) and XAROSA (EMSIS), respectively. Information regarding antibodies and other reagents is provided in Reagents and resources table.

#### Serial block face EM

Sample blocks from hippocampal CA3 regions of 0.5 x 0.5 mm were cut, mounted, and inserted into a Gatan 3View stage (Gatan) built in a Jeol JSM-7100F scanning electron microscope (Jeol). For imaging, the sample stage was biased with a 500V positive charge to account for sample charging during the scanning process. For the acquisition, 5×5 or 7×7 nm pixel size images were scanned, followed by the repeated ablation of 50 nm or 70 nm sections. The acquisition was controlled by the Gatan Digital 485. Micrograph software, which was also used for stack alignments. Further processing of the datasets was performed in Fiji. To analyse MFBs in the CA3 hippocampal region, subvolumes of such structures were randomly chosen and extracted to obtain smaller stacks. Images were contrast adjusted and, to reduce file size some acquisitions were adjusted to 8-bit images and scaled by a factor of 0.5. Afterwards, the volumes were rendered in Imaris v8/v9 (Bitplane; see Reagents and resources table) by manually identified membranes and outlining the structures in individual sections and creation of a 3D surface. All analysis steps (including selection of mossy fiber boutons) were performed by a researcher blinded to experimental group.

### Analysis

#### Analysis of antibody staining intensities

Intensity of ADAM10 immunosignal of stained MEF cells was analysed on background-subtracted widefield images using Fiji software [43] by an experimenter blind to the genotype. All cells within one image were outlined after thresholding, and average intensities measured.

#### Analysis of synaptic staining

ADAM10 presence at inhibitory and excitatory synapses was analysed using Fiji. Average projections of confocal images were used. VGLUT1 or VGAT was detected using the “find maxima” function in the full field of view excluding 1 µm edge. For each analysis 3 control ROIs per image were manually chosen within the dendrite but excluding synaptic spots (6 control ROIs in total per image). Average intensities in circular ROIs with 0.55 µm diameter were measured in all channels. For each image VGLUT1 and VGAT values were normalized to 1, ADAM10 was normalized so that all (inhibitory and excitatory) synapses average to 1. Changes in ADAM10 intensities at VGAT or VGLUT1 positive spots would be reflected in these normalized values. For quantification of the percentage of ADAM10 positive synapses, synapses were defined as ADAM10 positive if the ROIs contain more than 2 times the ADAM10 intensity than the average control ROIs within the dendrite.

#### Analysis of mossy fibers

Suprapyramidal mossy fiber and infrapyramidal mossy fiber were manually measured in images of sagittal sections in Fiji by a researcher blind to the genotype as described in [15].

#### Analysis of mossy fiber boutons in IHC

STED images of MFBs in cryo sections were analysed as follows: Munc13-1 and bassoon clusters were detected using the automated wavelet transform decomposition of the PySODA programme [57] based on [34] by a researcher blind to the genotype. Segmentation parameters were set to scales 3 and 4 as in [57]. Clusters of areas smaller than 17 pixels, or short axis of smaller than 4 pixels were discarded.

#### Statistical analysis and image representation

Statistical analysis was performed using Prism version 9 (GraphPad). 2-WAY repeated measures ANOVA p values for effect of genotype or treatment unless otherwise stated. Individual channels in multi-colour micrographs are contrasted for better representation, with set minimum and maximum identical if groups need to be compared using Fiji. No other modifications were done, unless otherwise stated. All analysis was performed on raw images. Figures were prepared using Adobe illustrator, based on graphs from Prism and images from Fiji.

## Results

### ADAM10 is targeted to the axonal compartment during neuronal differentiation and accumulates at presynaptic sites in adult neurons

Despite ADAM10’s importance in neuronal function, to date no systematic investigation of ADAM10 localization in different neuron types, during development, and at the subcellular level has been performed. We decided to address this question by immunofluorescence imaging in primary dissociated hippocampal cultures and brain sections as well as electron microscopy (EM). First, we thoroughly characterized an antibody directed against the intracellular carboxy (C)-terminus of ADAM10 (Fig 1a-d, S1a). High specificity of the anti-ADAM10 C-terminal antibody was confirmed using immunoblotting and immunocytochemistry (ICC) of wildtype (wt) and ADAM10 knockout (KO) mouse embryonic fibroblasts (MEF cells; [13], Fig. 1b, c), cortical extracts of cKO mice (Fig. 1d) and ICC of primary hippocampal rat neurons using a blocking peptide (Fig. S1a).

**Figure 1.**
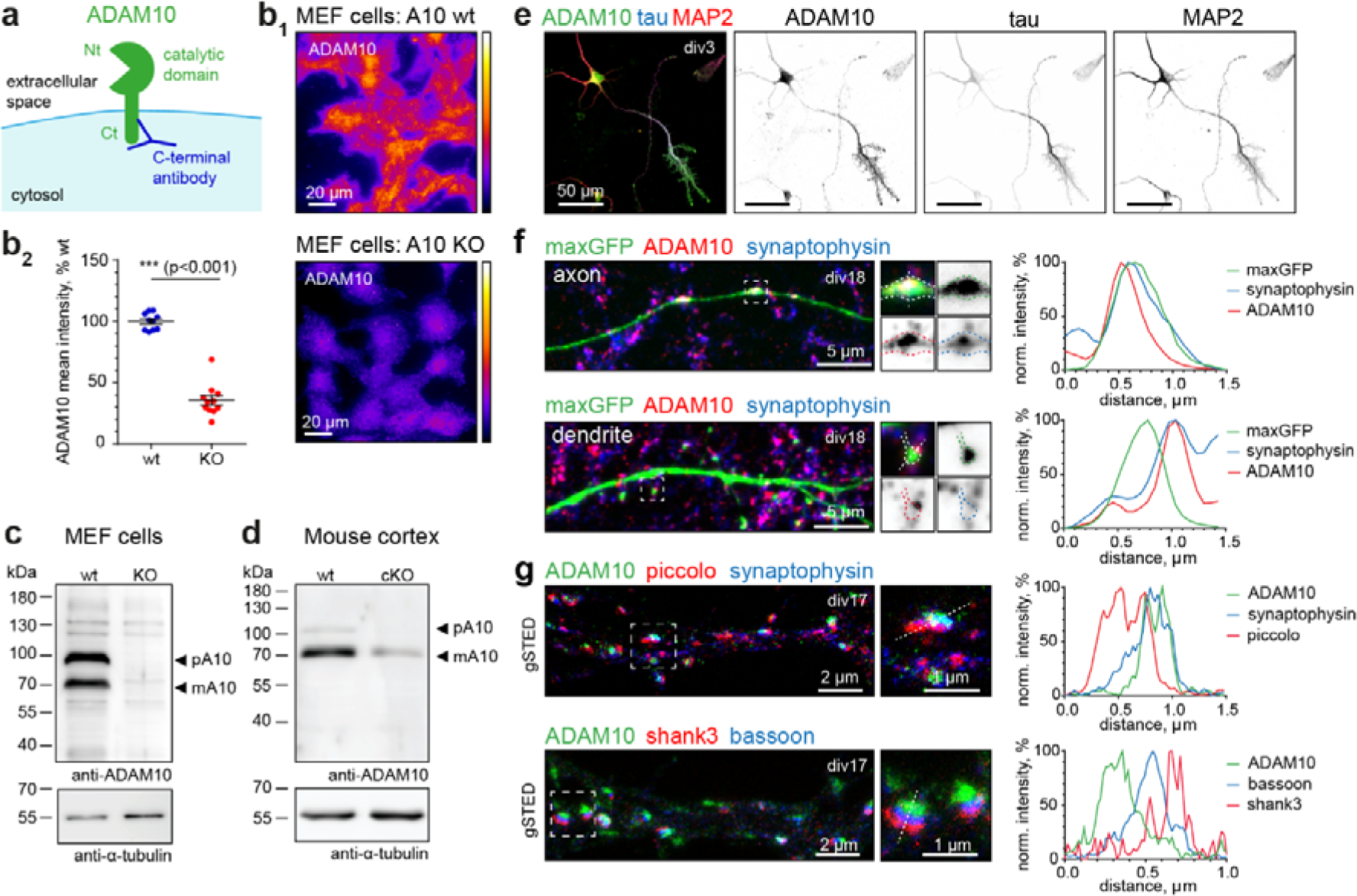
ADAM10 is strongly enriched at presynaptic sites **a)** Scheme of ADAM10 at the synaptic membrane with indicated C-terminal anti-ADAM10 antibody binding. Nt: N-terminus, Ct: C-terminus. **b-d)** Validation of the C-terminal ADAM10 antibody. **b)** Immunocytochemistry for ADAM10 in wildtype (+/+, wt) and knockout (-/-, KO) MEF cells. Representative widefield image **(b_1_)** and quantification **(b_2_)** show strong reduction of ADAM10 immunoreactivity in KO MEF cells. n=10 (wt), n=11(KO) images from 1 MEF cell preparation. 2-tailed unpaired Student‘s t-test. Data are represented as mean +/- SEM. **c)** Immunoblot analysis of ADAM10 wt and KO mouse embryonic fibroblast (MEF) cell lysates, detected with C-terminal ADAM10 antibody, indicates loss of ADAM10 bands corresponding to the precursor and mature form of the protease in KO cells. pA10: precursor of ADAM10; mA10: mature ADAM10. **d)** Immunoblot of P21 A10 cKO and wt cortical extracts shows strong reduction in the ADAM10 signal in the cKO. pA10: precursor of ADAM10; mA10: mature ADAM10. **e)** Representative maximum projections of confocal images of hippocampal primary cultures at div3. Immunostaining for ADAM10 (green), the axonal marker tau (blue), and MAP2 (red) as a dendritic marker. Note the strong enrichment of ADAM10 at the axon and axonal growth cones already in young cultures. **f)** Left: Representative maximum projection of confocal images of a div18 primary rat hippocampal neuron, transfected with a maxGFP cell fill (green) and stained for ADAM10 (red) and the presynaptic vesicle marker synaptophysin (blue) in an axon and at a dendrite. ADAM10 is present at presynaptic boutons. Note that dendritic spines are largely devoid of ADAM10. Right: Line scans of indicated axonal bouton and dendritic spine. **g)** Representative gated STED images of mature rat hippocampal primary neurons (div17) stained for ADAM10 (green), in combination with presynaptic cytomatrix of the active zone (CAZ) protein piccolo (red) and the vesicle marker synaptophysin (blue) or the presynaptic CAZ protein bassoon (blue), and the postsynaptic scaffold shank3 (red). Boxes indicate position of zoom-ins, lines were used for the line profiles shown. Note the localization of ADAM10 on the presynaptic (bassoon, blue) site. Right: Line scans of indicated synapses. See also **Figure S1.**

We observed ADAM10 staining not only in excitatory cells, but also in a variety of inhibitory neurons including GAD67, parvalbumin and calretinin-positive cells (Fig. S1b). We found that during neuronal differentiation ADAM10 can be detected already at day in vitro (div) 3 and, surprisingly, besides a somatodendritic expression, is strongly targeted to the axon (Fig. 1e). We next performed a series of ICCs of ADAM10 with different pre- and postsynaptic markers in mature rat hippocampal neuronal cultures (Fig. 1f,g, S1c-f). Since the ADAM10 antibody recognizes the C-terminal cytosolic domain rather than the extracellular N-terminus, it is well-suited for determining the pre- or postsynaptic localization of the protein. To map the subcellular localization of ADAM10 we examined neurons filled with maxGFP. Confocal imaging revealed a punctate distribution of ADAM10 throughout the neuron (Fig. S1a), and the fluorescence signal was closely associated with the presynaptic vesicle marker synaptophysin at axonal boutons (Fig. 1f, upper panel). When looking at dendritic spines visualized by maxGFP, ADAM10 was predominantly located near the edges, once again closely associated with synaptophysin (Fig. 1f, lower panel), showing surprisingly little signal in dendrites and postsynaptic sites. To verify the presynaptic localization of ADAM10, we turned to super-resolution imaging and performed gated stimulated emission depletion nanoscopy (gSTED). We observed that ADAM10 is present mainly in the presynaptic compartment of synapses. It colocalizes with the presynaptic markers bassoon and piccolo and not the postsynaptic marker shank3 (Fig. 1g).

We next investigated the presence of ADAM10 at different types of presynaptic terminals (Fig. S1c-f). Mature primary hippocampal cultures stained for ADAM10 and the presynaptic vesicle markers VGAT (inhibitory synapse) and VGLUT1 (excitatory synapse) show colocalization of both markers with ADAM10 (Fig. S1c). Quantification revealed that approximately 80 % of inhibitory and excitatory synapses are ADAM10-positive and there is a correlation between ADAM10 and VGAT/VGLUT1 intensity (Fig. S1d,e, see methods). Line scans of confocal and gSTED images confirmed the localization of ADAM10 with both inhibitory and excitatory presynaptic vesicle markers (Fig. S1f). This is particularly intriguing as the presynaptic localization of ADAM10 implies potential unexplored functions of the protease.

### ADAM10 is present in presynaptic vesicles at hippocampal mossy fiber synapses

Given that primary hippocampal cultures comprise a variety of neuronal subtypes, we next set out to analyze the localization of ADAM10 in the hippocampus where its proteolysis-related functions have been previously explored [17, 36, 50]. Following diaminobenzidine (DAB) staining of a mouse brain we found a strong enrichment of ADAM10 in MFs, which arise from dentate granule neurons and connect the dental gyrus with the CA3 principal neurons (Fig. 2a). Mossy fiber boutons carry specialized excitatory synapses that contact postsynaptic thorny excrescences on proximal dendrites. A single MFB can contain more than a dozen release sites onto one CA3 neuron [39]. A single dentate gyrus (DG) granule cell forms on the order of a dozen MFBs connecting with different postsynaptic cells in the CA3 [3]. As these contacts are vital for hippocampal function and information processing, we decided to investigate the exact localization and function of ADAM10 at MFs in more detail. EM of DAB-labelled sections revealed ADAM10 staining in the presynaptic MFBs (Fig. 2b, pink arrows) opposing the postsynaptic densities of CA3 thorny excrescences, that was not found in the anti-ADAM10 antibody-lacking control (Fig. 2b, yellow arrows). Immunogold labeling with the same antibody indicated that ADAM10 localized to synaptic vesicles (Fig. 2c). Gold particles were exclusively located on the outside (cytosolic side) of vesicles, as would be expected if ADAM10 is integrated into the vesicular membrane (Fig. 2c). Of note, we did not observe gold particles associated with the presynaptic membrane, suggesting that most presynaptic ADAM10 is associated with synaptic vesicles.

**Figure 2.**
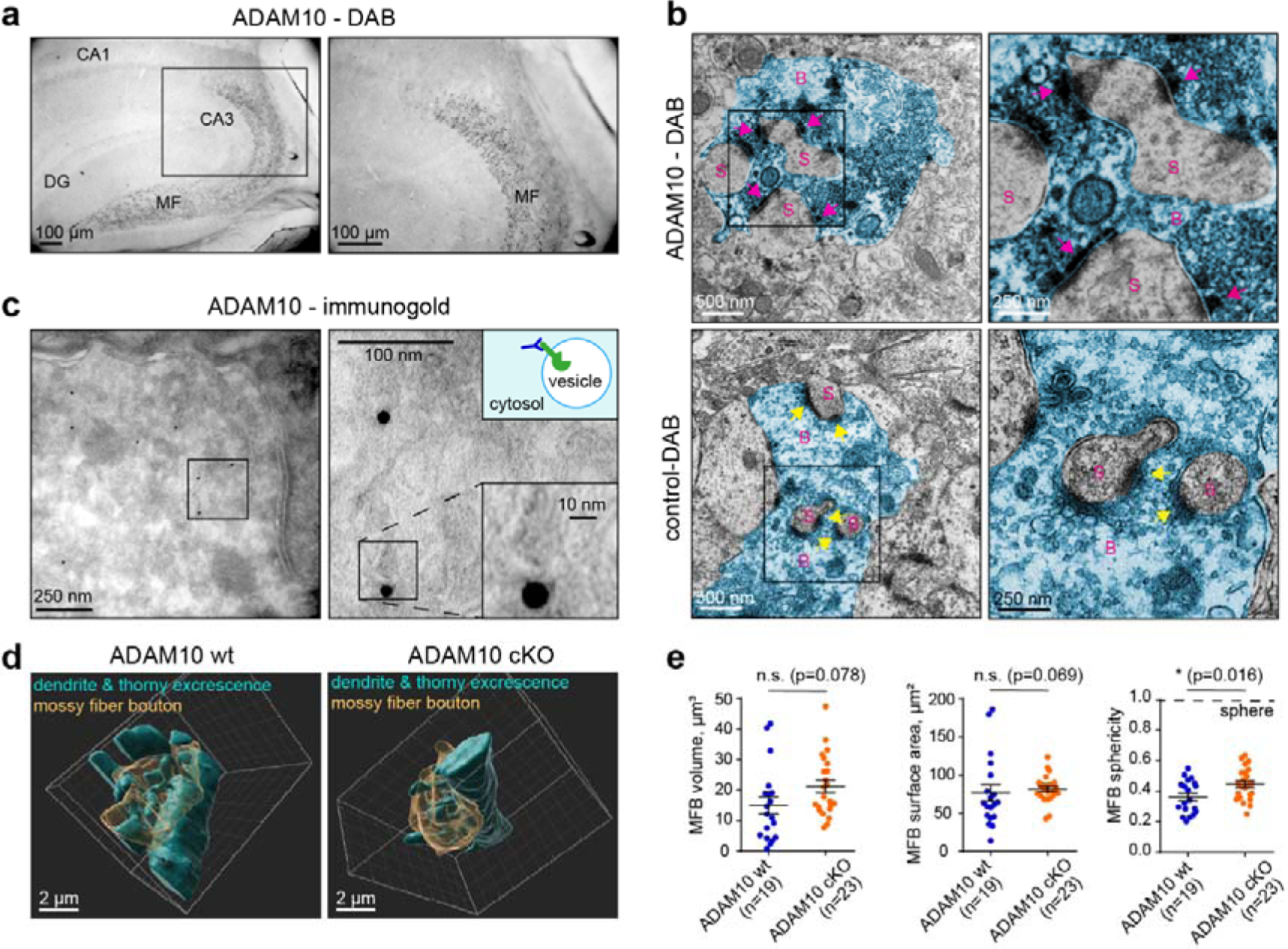
ADAM10 in enriched in vesicles of mossy fiber boutons, that show only minor morphological changes in cKO animals **a)** DAB staining of ADAM10 in an adult wildtype mouse hippocampus shows strong enrichment of ADAM10 in mossy fibers. DG: dentate gyrus. MF: Mossy fibers. **b)** High magnification of ADAM10 DAB and control (without primary antibody) staining in MF-CA3 synapses. Note the strong ADAM10 localization to the presynaptic site (pink arrow) and lack of DAB staining at the presynaptic membrane in the control (yellow arrowhead). S: dendritic spine. B: mossy fiber bouton (false coloured in blue). **c)** Immunogold EM of a P21 wildtype mouse with focus on hippocampal mossy fiber boutons. Note that lack of gold particles at the synaptic membrane and that gold particles localize to the outside of vesicles, as the antibody detects ADAM10’s cytosolic C-terminus (see the scheme). **d)** Example of ADAM10 cKO and wt mossy fiber bouton 3D reconstructions from SBEM data. See also **Videos S1-S4**. **e)** Quantification of MFB volume, surface area and sphericity. n=19 (wt), n=23 (cKO) boutons from 3 animals each. 2-tailed unpaired Student‘s t-test (volume, surface area) and Mann Whitney U-test (sphericity). Data are represented as mean +/- SEM. See also **Figure S2**.

The strong enrichment of ADAM10 in MFBs suggests a prominent role at this synapse. As there are no reports related to ADAM10 function at MFBs, we decided to investigate this in more detail using ADAM10 cKO mice. This mouse model lacks ADAM10 in excitatory neurons of the forebrain through CaMKII-driven Cre recombinase expression, this also includes dentate gyrus granular cells carrying MFBs [36]. An anatomical examination using the MF marker synaptoporin and the nuclear marker DAPI showed no alterations in the overall morphology of intra- and suprapyramidal mossy fiber tracts (length ratio, Fig. S2a). This suggests that hippocampal connectivity in ADAM10 cKO mice remains normal.

To investigate the effect of ADAM10 on MFB morphology in greater detail we then used serial block face EM (SBEM) to reconstruct 19 control and 23 ADAM10 cKO MFBs in three dimensions (Fig. 2d, Videos S1-4). No differences were observed in the volume or surface area of the MFBs although the MFBs of ADAM10 cKO mice were simpler, as indicated by sphericity measurements (Fig. 2f). Next, for the analysis of presynaptic active zones in MFBs, we performed immunohistochemistry (IHC) and STED imaging in cryosections of ADAM10 wildtype and cKO mice (Fig. S2b). We used bassoon as a marker for presynaptic active zones and munc13-1, which, due to its important function in vesicle priming, is a good proxy for the ready releasable pool of vesicles [22, 41]. These markers label multiple release sites within the same MFB which are distinguishable by STED imaging [7]. There was only a minor increase in average bassoon cluster sizes and no changes in munc13 clusters, suggesting ADAM10 may not affect vesicle priming (Fig. S2c). Taken together, these data indicate that neuron-specific deletion of ADAM10 has a minor impact on the ultrastructure of presynaptic contact sites and MFB morphology. This is surprising because the deletion of ADAM10 in the CA1 region of the hippocampus has a profound effect on synaptic morphology [36] and suggests that the protease may have independent functions in MF synapses.

### Loss of ADAM10 diminishes DG-CA3 mossy fiber synaptic facilitation

As ADAM10 is highly enriched at synaptic vesicles in MFBs, we reasoned that it may have an impact on transmitter release. To test this, mossy fibers were stimulated, and field excitatory postsynaptic potentials (fEPSP) were recorded in the CA3 stratum radiatum (Fig. 3a-d, S3a-d). The mossy fiber origin of the fEPSPs was confirmed at the end of every experiment by the subsequent loss of response upon applying the group II mGluR agonist DCGIV, which selectively inhibits presynaptic release at DG-MF synapses but not at associational or commissural synapses [Fig. 3c, [23]]. There were no differences in stimulation threshold or fiber volley amplitude in slices from WT and ADAM10 cKO mice, suggesting that ADAM10 does not change excitability of DG granule cells (Fig. S3a,b).

**Figure 3.**
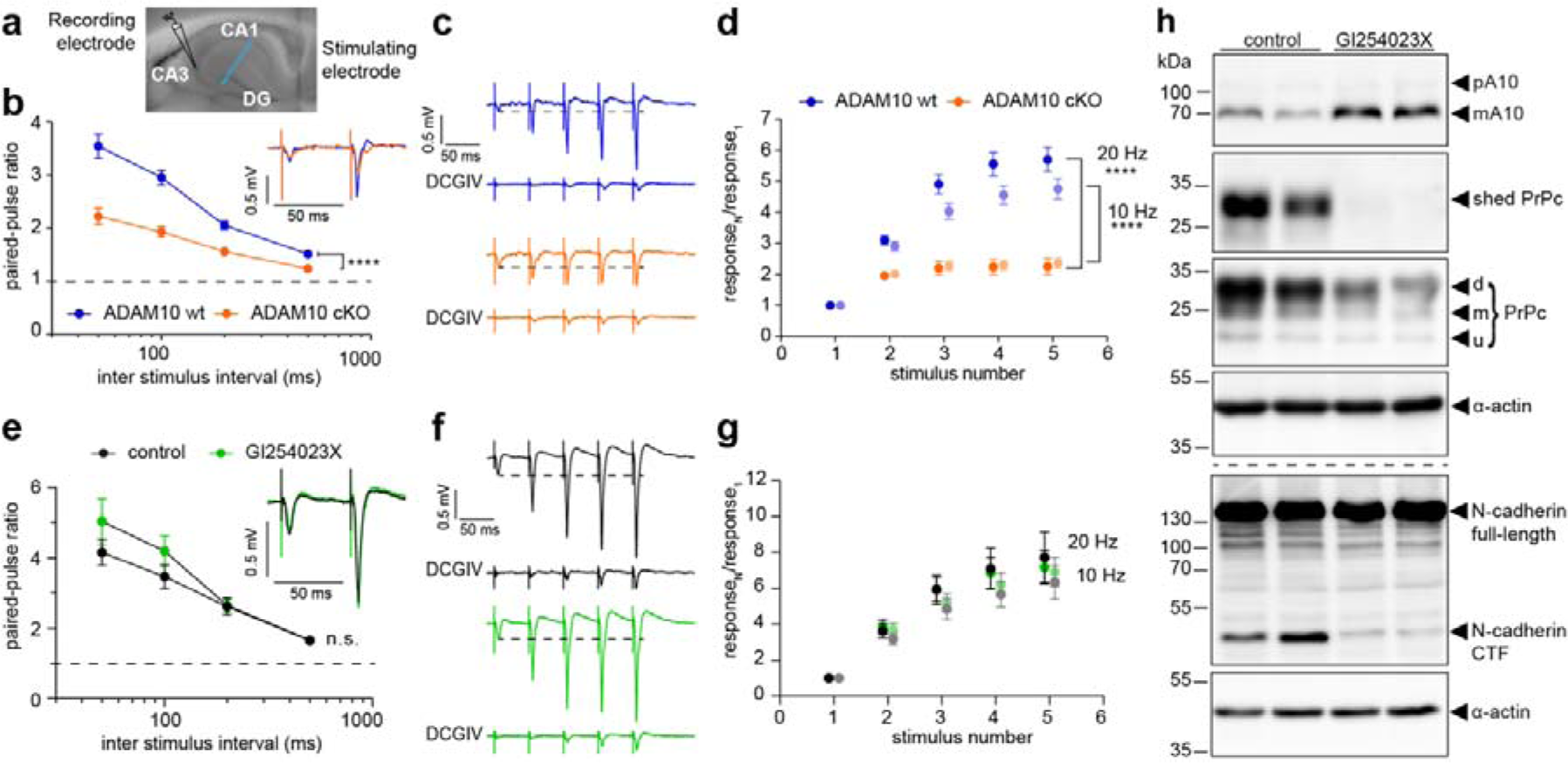
ADAM10 is required for the expression of presynaptic mossy fiber short-term plasticity which does not depend on the enzymatic activity of the protease **a)** Image of an acute hippocampal slice with indicated positions for stimulating (MF) and recording (CA3) electrodes. **b-d)** Mossy fiber plasticity of ADAM10 wt and cKO animals. **b)** Paired pulse facilitation ratio at different stimulation frequencies and example traces (average of 5 sweeps, inlet) of evoked fEPSPs at 20 Hz of ADAM10 cKO and wt mice. ADAM10 cKO show impaired facilitation. n= 18 (wt), n= 19 (cKO) slices from 5 mice each. Two-way repeated measures ANOVA. ****p < 0.0001. Data are represented as mean +/- SEM. **c)** Example traces of train facilitation at 20 Hz in wt and cKO slices. Application of the group II mGluR agonist DCGIV (1 µM) leads to a loss of response and is used to prove the mossy fiber origin of the detected signal. **d)** Quantification of the ratio calculated from the fEPSP amplitudes measured in response to train facilitation. ADAM10 cKO slices show an impaired response to train stimulation at 20 Hz (dark colours) and 10 Hz (light colours). 2-way repeated measures ANOVA. **** p<0.0001. n=18 slices (wt), n=19 slices (cKO) from 5 mice each. Data are represented as mean +/-SEM. **e-h)** MF plasticity of wt animals with or without ADAM10 inhibitor (GI254023X) treatment. **e)** Paired pulse facilitation ratio at different stimulation frequencies and example traces [average of 3 (GI254023X) or 4 (ctr) sweeps, inlet] of evoked fEPSPs at 20 Hz of wt mouse slices upon inhibition of ADAM10 activity. Application of the ADAM10 inhibitor GI254023X does not affect synaptic facilitation. Two-way repeated measures ANOVA. Treatment p = 0.3113. n = 12 slices (control); n = 11 slices (GI254023X) from 3 mice each. Data are represented as mean +/-SEM. **f)** Example traces of train facilitation at 20 Hz in wt slices with and without ADAM10 inhibitor (GI254023X). Application of the group II mGluR agonist DCGIV (1 µM) leads to a loss of response and is used to prove the mossy fiber origin of the detected signal. **g)** Quantification of the ratio calculated from the fEPSP amplitudes measured in response to train facilitation. Treatment with ADAM10 inhibitor does not change the ratio calculated in response to train stimulation at 20 Hz (dark colours) and 10 Hz (light colours). Two-way repeated measures ANOVA. p=0.5764 (10 Hz), p=0.9124 (20 Hz). n = 12 (control); n = 11 slices (GI254023X) from 3 mice each. Data are represented as mean +/-SEM. **h)** Immuno blot analysis of acute hippocampal wt slices untreated or treated with GI254023X confirming that the application of the ADAM10 inhibitor does in fact reduce ADAM10 activity. Note the reduced substrates cleavage (PrPc to shed PrPc, N-cadherin to C-terminal fragment CTF) in the GI254023X group. d: di-glycosylated, m: mono-glycosylated, u: unglycosylated See also **Figure S3.**

A characteristic feature of DG-CA3 mossy fiber synapses is the pronounced frequency dependent facilitation of transmitter release. This property is mediated by presynaptic mechanisms [reviewed in [14, 33, 37]]. Importantly, we found that in ADAM10 cKOs paired-pulse facilitation is significantly reduced over a wide range of interstimulus intervals (ISIs, Fig. 3b) while initial responses were similar (Fig S3c). Facilitation evoked by short trains of stimuli was also impaired for frequencies of 10 and 20 Hz (Fig. 3c,d, S3d). Thus, loss of ADAM10 perturbs presynaptic facilitation without affecting synaptic morphology or responses to single stimuli.

### ADAM10 effects on presynaptic plasticity are independent of its proteolytic activity

Taking into account the multiple synaptic substrates of ADAM10 we next asked if the ADAM10 proteolytic activity is necessary for presynaptic facilitation. To test this, we inhibited the enzymatic activity of ADAM10 by pre-incubation of slices with the selective ADAM10 inhibitor GI254023X [16] three hours prior to the electrophysiological experiments (Fig. 3e-h, S3e-h). Surprisingly, GI254023X had no effect on facilitation either induced by paired pulses or trains (Fig. 3e-g, S3e-h). The amplitude of the first response was also unchanged (Fig. S3g). Importantly, we collected the slices after the experiments and verified the inhibition of ADAM10 activity after GI254023X treatment (Fig. 3h). The specific ADAM10 cleavage products shed prion protein (sPrPc; [28]) and N-cadherin C-terminal fragment [38] were strongly reduced in GI254023X-treated slices (Fig. 3h). Thus, the presence of ADAM10 protein per se rather than the enzymatic activity is important for its role in MF short-term plasticity.

### ADAM10 is involved in synaptotagmin 7-dependent mechanism of presynaptic facilitation

In search for a molecular explanation of how ADAM10 contributes to presynaptic plasticity of MFBs, we noted the decreased MF short-term plasticity in the absence of ADAM10 closely resembles the reduced facilitation previously observed at this synapse in syt7 KO mice [20]. Syt7 is a transmembrane calcium sensor protein. It contains two cytosolic C2 domains, that allow Ca^2+^ binding, as well as binding to phospholipids and other interaction partners. Among synaptotagmins, syt7 has one of the highest Ca^2+^ affinities and slow Ca^2+^-un/binding kinetics as well as slow phospholipid disassembly rates [reviewed in [18, 19]].

We were therefore interested to study if ADAM10 and syt7 play an orchestrated function in MF short-term plasticity (Fig. 4,5). Immunoblot analysis revealed a 20 % reduction in syt7 levels in hippocampal slices from ADAM10 cKO mice. In contrast, the major calcium buffer in MFs, calbindin was unchanged suggesting importantly that calcium buffering is unaffected by the loss of ADAM10 (Fig. 4a). Similarly, the synaptic vesicle-associated protein VAMP1 was also not changed (Fig. 4a) confirming observations by Cozzolino et al., indicating that presynaptic vesicle number is not affected in cKO ADAM10 mice [10]. We reasoned that this reduction in syt7 protein levels could not alone account for the profound loss of facilitation. Therefore, we next asked whether ADAM10 and syt7 can be found in the same complex. For this we performed an endogenous immunoprecipitation of ADAM10 from synaptosomes of wildtype mice. Indeed, we found that syt7 co-precipitated with ADAM10, irrespectively of Ca^2+^ presence (Fig. 4b). This finding was further confirmed by heterologous co-immunoprecipitations from Neuro-2a (N2a) cells (Fig. 4c).

**Figure 4.**
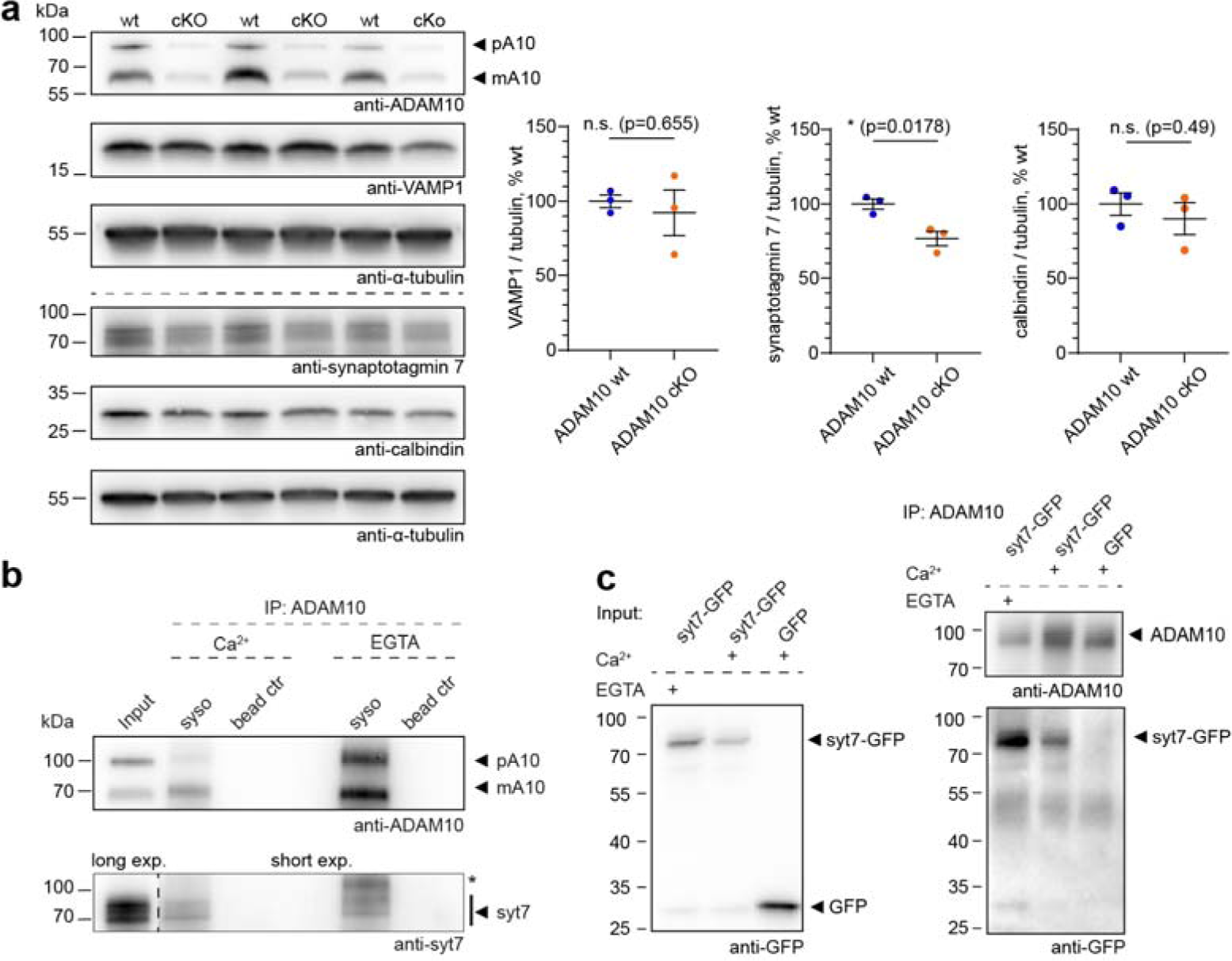
ADAM10 acts via the syt7 pathway **a)** Syt7 levels in hippocampus of ADAM10 cKO mice are reduced, while the major mossy fiber calcium buffer calbindin and the vesicle marker VAMP are unchanged. n=3 acute slice preparations of 3 animals (same slices as in Figure 4). Unpaired, 2 -tailed Student’s test. **b)** Synaptic syt7 associates with ADAM10 in a Ca^2+^-independent manner. Endogenous Co-immunoprecipitations from mouse synaptosomes in presence of 200 µM CaCl_2_ or 2 mM EGTA. * unspecific band. Note the different exposure times for the lower blot. syso: synaptosomes; ctr: control. **C)** Heterologous Co-immunoprecipitations from Neuro-2a cells shows syt7-GFP is in one complex with ADAM10 in both calcium (200 µM) and calcium-free (2mM EGTA) conditions.

**Figure 5.**
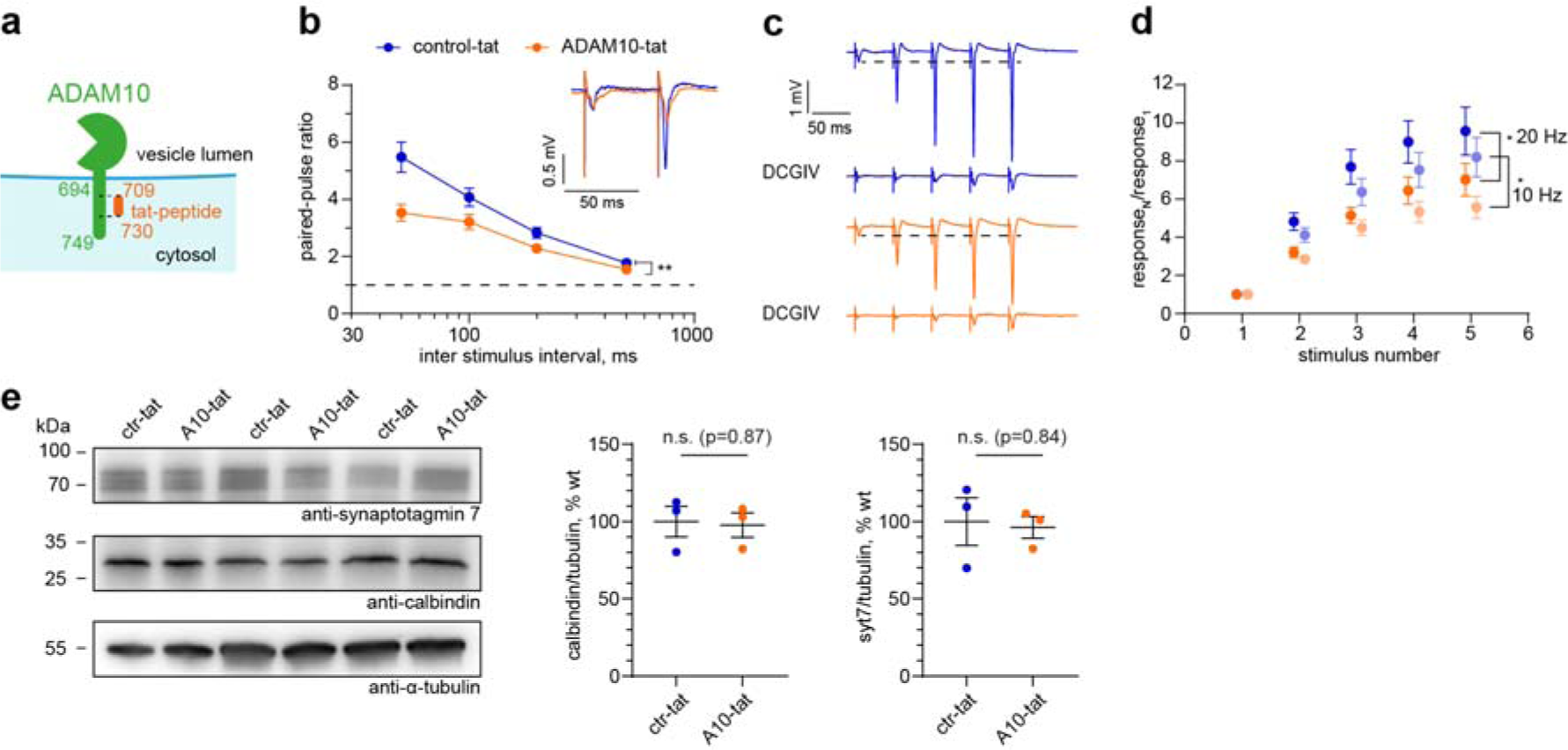
ADAM10 C-terminus is required for mossy fiber short-term plasticity **a)** Scheme of tat-peptide in relation to ADAM10 C-terminus. l: length, aa: amino acids. **b)** Paired pulse facilitation ratio at different stimulation frequencies and example traces (average of 5 sweeps, inlet) of evoked fEPSPs at 20 Hz in hippocampal slices treated with ctr-tat or ADAM10-tat peptide. Application of ADAM10 C-terminus targeted tat-peptide leads to impairment in facilitation. Two-way repeated measures ANOVA. **p = 0.0086. n = 12 slices from 3 mice each. Data are represented as mean +/-SEM. **c)** Example traces of train facilitation at 20 Hz in both experimental groups. Application of the group II mGluR agonist DCGIV (1 µM) is used as control for the mossy fiber origin of the detected signals. **d)** Quantification of the ratio calculated from the fEPSP amplitudes measured in response to train facilitation. Impaired response upon ADAM10-tat-peptide treatment to train stimulation at 20 Hz (dark colours) and 10 Hz (light colours). 2-Way repeated measures ANOVA. * p=0.0294 (10 Hz), * p= 0.0409 (20 Hz). n=12 slices from 3 mice each. Data are represented as mean +/-SEM. **e)** Syt7 levels are not changed in hippocampal slices after treatment with ADAM10-tat peptide compared to control-tat peptide. n=3 acute slice preparations of 3 animals. Unpaired, 2 -tailed Student’s test. See also **Figure S4.**

Since both ADAM10 and syt7 are present on the vesicular and plasma membrane, respectively, association in one complex would have to occur via the 55 amino acid long cytosolic tail of ADAM10 (scheme Fig. 5a). Notably, the C-terminus of ADAM10 contains two proline-rich PxxP motifs that are known to be sites of protein-protein interaction [11]. To perturb the association with potential binding partners, we therefore generated a cell-permeable tat-peptide corresponding to mouse ADAM10 amino acids 709-730 covering all PxxP motifs (ADAM10-tat, Fig. 5a). An almost identical tat-peptide was previously shown to disturb C-terminal interactions of ADAM10 [30]. As a control peptide we substituted alanines for all prolines. Based on the electrophysiology and co-immunoprecipitation results, we hypothesized that an association between the C-terminal domain of ADAM10 and syt7 is required for MF facilitation. We performed a new set of electrophysiological experiments, this time using slices of wildtype mice in the presence of either ADAM10-tat or control-tat-peptide (Fig. 5b-d, S4). Basal transmission was unaffected by including 3 µM of ADAM10-tat-peptide compared to control-tat-peptide (Fig. S4a,b). However, application of ADAM10-tat-peptide but not control-tat-peptide reduced the paired-pulse ratio (Fig. 5b, S4c) and train facilitation (Fig. 5c,d, S4d), similar to the ADAM10 cKO phenotype (Fig. 4). Importantly, immunoblot analysis revealed that ADAM10-tat-peptide does not decrease MF-CA3 facilitation by decreasing syt7 protein levels (Fig. 5e). The fact that applying the tat-peptide acutely mimics ADAM10 cKO strongly suggests that primarily the syt7-ADAM10 interaction is required for presynaptic facilitation. From these data we conclude that ADAM10 and syt7, in concert with other -to be identified-presynaptic proteins, cooperate to regulate MF short-term plasticity.

## Discussion

Considering that the Alzheimer Disease related alpha-secretase ADAM10 is a prominent disease-associated protease whose function has been intensively investigated from a substrate point of view, little was known about the cellular and subcellular localization of the protein in the brain. In this study, we show that ADAM10 displays a prominent axonal localization already very early in neuronal differentiation. During synaptogenesis ADAM10 accumulates at presynaptic sites. In the hippocampus ADAM10 is highly enriched in MFBs of dentate granule cells where it is almost exclusively found on presynaptic vesicles. By combining functional and biochemical studies we discover that ADAM10, in concert with syt7, increases facilitation at this synapse independently of its proteolytic activity. This sheds light on previously undiscovered physiological functions of the protease.

Previous studies have focused on the proteolytic function of ADAM10 and the identification of substrates in the brain. As several of the most important substrates including APP, prion protein and neuroligins are expressed postsynaptically, it was largely assumed that ADAM10 would be mainly expressed in dendrites and postsynaptic sites [40]. Of note, no high-resolution microscopy studies with knock out validated antibodies were available in the literature, leaving the subcellular localization of ADAM10 still an open question. Indeed, several prominent ADAM10 substrates are not exclusively postsynaptic. APP, for example, is also enriched at the presynapse [8, 56]. Other important ADAM10 substrates, such as the Notch receptor, low density lipoprotein receptor, neurexin-1 and neuronal cell adhesion molecule, are also presynaptic [13, 26, 50, 60]. As the proteolytic domain is extracellular, either a predominantly presynaptic or postsynaptic localization does not preclude shedding of the extracellular domains of proteins located in adjacent membranes. Consistent with a predominantly presynaptic localization is the finding that ADAM10 is enriched in the purified brain vesicle fraction [29]. Likewise, a mass spectrometry profiling of the ADAM10 complex indicated the presence of several active zone components including piccolo and bassoon [10]. These observations support our data on the presynaptic localization of ADAM10.

Our results also explain a previous observation on activity-dependent substrate shedding by ADAM10: shedding of neuroligin 3, which promotes glioma growth, is neuronal activity dependent and is prevented by inhibiting action potential-dependent transmitter release with tetrodotoxin [53]. In light of our findings, we can hypothesize how this activity dependence of ADAM10-mediated shedding could occur. When ADAM10-decorated vesicles fuse and transmitter is released, the catalytic domain, which was previously inside the vesicles, will now be exposed extracellularly within the synaptic cleft where it can cleave its substrates. We did not detect ADAM10 immunogold labeling of synaptic or other surface membranes in CA3 of the hippocampus, suggesting that in resting conditions little ADAM10 is present on the surface of neurons. It would be interesting to investigate in the future if this hypothesis holds true.

In hippocampal sections, the accumulation of ADAM10 at MF-CA3 synapses is striking. These synapses have a unique morphology, molecular composition and exhibit strong frequency facilitation contributing to their designation as a ‘detonator’ synapse [4, 25]. The conditional deletion of ADAM10 had no major impact on DG-CA3 connectivity or the morphology of MFBs. We also did not observe a change in munc13-1 clusters, which suggests that ADAM10 does not affect the probability of presynaptic vesicle release [41]. This suggestion is supported by our observation that there is no change in the amplitude of the EPSPs recorded in response to the first stimuli in ADAM10 cKO mice compared to wt. Instead, ADAM10 promotes frequency facilitation, a hallmark of this synapse [42, 49]. Facilitation is high at synapses where the probability of transmitter release after single action potentials is low. At such synapses, single action potentials will not cause postsynaptic neurons to spike. Instead, a burst of action potentials in the presynaptic neuron is required before the postsynaptic neurons will spike and transmit information within a circuit. As ADAM10 promotes facilitation at the MF-CA3 synapse, loss of ADAM10 is expected to decrease information transmission from the dentate gyrus to the hippocampus, thus contributing to the cognitive and learning deficits associated with loss of ADAM10 [36].

Unexpectedly, the impairment of MF facilitation in cKO mice is not a result of loss of ADAM10 proteolytic activity as pharmacological inhibition of ADAM10 in wildtype slices had no impact in this plasticity paradigm. Importantly, the application of a tat-peptide, mimicking the cytosolic tail of ADAM10, replicates the observed physiological phenotype of ADAM10 cKO mice. This implies that the identified plasticity defects result from the involvement of ADAM10’s cytosolic C-terminus and are independent of ADAM10 extracellular enzymatic activity. Furthermore, this finding indicates that the observed plasticity defects are not attributable to the minor morphological changes observed in MFBs of the cKO mice. Our results underscore the essential role of cytosolic protein interactions, rather than the proteolytic function, of ADAM10 in a paradigm of plasticity, highlighting the diverse cellular functions carried out by the same protein. These findings also align with the importance of the cytosolic tail in ADAM10 basolateral localization in epithelial cells [58] and in postsynaptic localization via interaction with the scaffolding protein SAP97 [30]. Of note, a closely related protease ADAM17, which is also expressed in the brain and shares some substrates with ADAM10 [40, 61], does not seem to compensate for the structural role of ADAM10. Our data reveal a hitherto poorly recognized non-proteolytic role of ADAM10 in presynaptic vesicle release. Therapeutically targeting ADAM10 by small molecules or other means may therefore also cause an altered synaptic plasticity possibly associated with unwanted side effects.

What could be the molecular underprints explaining the ADAM10 cKO facilitation deficits? There is not much known about presynaptic interaction partners of ADAM10. Recently, a cytomatrix of the active zone (CAZ) protein, piccolo, was reported to associate with ADAM10 [10]. However, neither piccolo knockout animals, nor neurons double deficient for piccolo and the highly homologous protein bassoon, show presynaptic facilitation deficiencies [1, 9, 32]. Interestingly, at most strongly facilitating synapses, including the hippocampal MFBs, the calcium sensor syt7 is required [19]. In contrast to the fast sensors that mediate rapid neurotransmitter release, such as vesicular syt1, syt7 localizes to the presynaptic membrane and is a higher affinity calcium sensor with slow on/off kinetics [45, 54]. In syt7 knockout mice facilitation is abolished, whereas the initial probability of release and presynaptic residual calcium concentrations are unaltered [20]. Whereas ADAM10 cKO does not abolish facilitation, it strongly reduces it without apparent effects on the initial responses. Indeed, we found that syt7 and ADAM10 interact suggesting that ADAM10 exerts it effects on facilitation via syt7. Loss of ADAM10 reduced hippocampal levels of syt7 by about 20 %, which may contribute to the reduction in EPSP facilitation. However, acutely interfering with cytosolic protein-protein interactions with the ADAM10-tat-peptide similarly decreased facilitation suggesting a more direct role of ADAM10 in maintaining frequency facilitation. Our data support the conclusion that ADAM10 and syt7 are present in one complex which is formed irrespectively of calcium presence. As ADAM10 and syt7 are associated with vesicles and the plasma membrane respectively, several scenarios are possible: 1) ADAM10/syt7 interaction may participate in vesicle docking, 2) ADAM10 may recruit syt7 to presynaptic release sites with docked vesicles, 3) ADAM10 may stabilize syt7 complexes. Future research will shed light on the mechanistic aspects of ADAM10 and syt7 interplay for the control of presynaptic short-term plasticity. Whether ADAM10 also plays a role in vesicle release and facilitation at other synapses or only in select synapses remains an open question. Syt7 is diffusely present along axonal and dendritic membranes where it is involved in exocytosis of other vesicle types, such as lysosomes or dense core vesicles [35, 52]. Since both proteins are also expressed in secretory cells [6, 48], it will be interesting to investigate if ADAM10 also plays a role in non-action potential driven vesicular release.

## Abbreviations

aCSF: artificial cerebrospinal fluid solution,
AD: Alzheimer’s disease,
ADAM: A Disintegrin And Metalloprotease,
APP: amyloid precursor protein,
BB: blocking buffer,
BSA: bovine serum albumin,
CAZ: cytomatrix of the active zone,
cKO: conditional KO,
C-terminus: carboxy-terminus,
DG: dentate gyrus,
div: day in vitro,
EM: electron microscopy,
fEPSP: field excitatory postsynaptic potentials,
gSTED: gated STED,
HD: Huntington’s disease,
HRP: horse reddish peroxidase,
ICC: immunocytochemistry,
IHC: immunohistochemistry,
KO: knockout,
MEF: mouse embryonic fibroblast,
MF: mossy fiber,
MFB: MF bouton,
N2a: Neuro-2a,
PB: phosphate buffer,
PPR: Paired-pulse ratio,
SBEM: serial block face EM,
sPrPc: shed prion protein,
STED: stimulated emission depletion,
syt: synaptotagmin,
TBS: tris buffered saline,
wt: wildtype.

## Acknowledgements

We would like to thank Sabine Graf and Jan Schröder (ZMNH, Hamburg) for help with electrophysiological recordings at the synchro slicer, Dr. Oliver Kobler (Combinatorial Neuroimaging Core Facility (CNI), LIN Magdeburg) for help with STED imaging, Dr. Antonio Virgilio Failla (UKE Microscopy Imaging Facility (UMIF)) for access and use of their Leica SP5 confocal microscope, Dr. Rudolph Reimer (Heinrich-Pette-Institut, Leibniz Institute for Experimental Virology, Hamburg) for help with SBEM data acquisition and microscope access and Abberior GmbH for access to the STEDYCON system. We thank Hermann Altmeppen for the anti-shed Prion antibody and Volker Haucke for the syt7-GFP construct. We acknowledge Marlies Rusch, Chudamani Raithore and Lisa Mallis for technical assistance and Theresa Felix for help with initial EM analysis. This research is supported by the German research foundation (CRC877 B12 to MM and A3 / Z3 to PS, Excellence Strategy – EXC-2049– 390688087, FOR2419 P2 and CRC1315 A10 to MM and FOR2419 P7 to CEG).

## Statements & Declarations

### Author Contributions

Designed experiments (JB, TF, CEG, PS, MM), performed experiments (JB, TF, CCR, LS, MG, MS, MM), analyzed and interpreted data (JB, TF, MB, JRB, FD, PS, MM), developed the manuscript concept and wrote the paper (JB, PS, MM), reviewed and edited the paper (JB, TF, CCR, CEG, MB, PS, MM) and funding acquisition (PS, CEG, MM).

### Data Availability

Materials (ADAM10 cKO mice) can be requested from Paul Saftig.

### Ethics approval

Animal experiments were carried out in accordance with the European Communities Council Directive (2010/63/EU) and the Animal Welfare Law of the Federal Republic of Germany (Tierschutzgesetz der Bundesrepublik Deutschland, TierSchG) approved by the local authorities of Schleswig-Holstein/Germany (V242-20464/2021 (31-4/21)) or of the city-state Hamburg (Behörde für Gesundheit und Verbraucherschutz, Fachbereich Veterinärwesen, from 21.04.2015, ORG781 and 1035) and the animal care committee of the University Medical Center Hamburg-Eppendorf.

