## Supplemental figures for "Non-canonical function of ADAM10 in presynaptic plasticity"

### Cellular and Molecular Life Sciences

**Julia Bär[1, 2], Tomas Fanutza[1, 2], Christopher C. Reimann[2], Lisa Seipold[3], Maja Grohe[3], Janike Rabea Bolter[1], Flemming Delfs[2], Michael Bucher[2], Christine E Gee[4], Michaela Schweizer[5], Paul Saftig[3]*, Marina Mikhaylova[1,2,6,*]**

[1] AG Optobiology, Institute of Biology, Humboldt Universität zu Berlin, Berlin, 10115 Germany

[2] Guest Group "Neuronal Protein Transport", Center for Molecular Neurobiology, University Medical Center Hamburg-Eppendorf, Hamburg, 20251 Germany; discontinued in August 2023

[3] Biochemisches Institut, Christian Albrechts-Universität Kiel, Kiel, 24098 Germany

[4] Department of Synaptic Physiology, Center for Molecular Neurobiology, ZMNH, University Medical Center Hamburg-Eppendorf, 20251 Hamburg, Germany

[5] Morphology and Electron Microscopy, University Medical Center Hamburg-Eppendorf, Center for Molecular Neurobiology, ZMNH, Hamburg, 20251 Germany

### Supplemental Figures

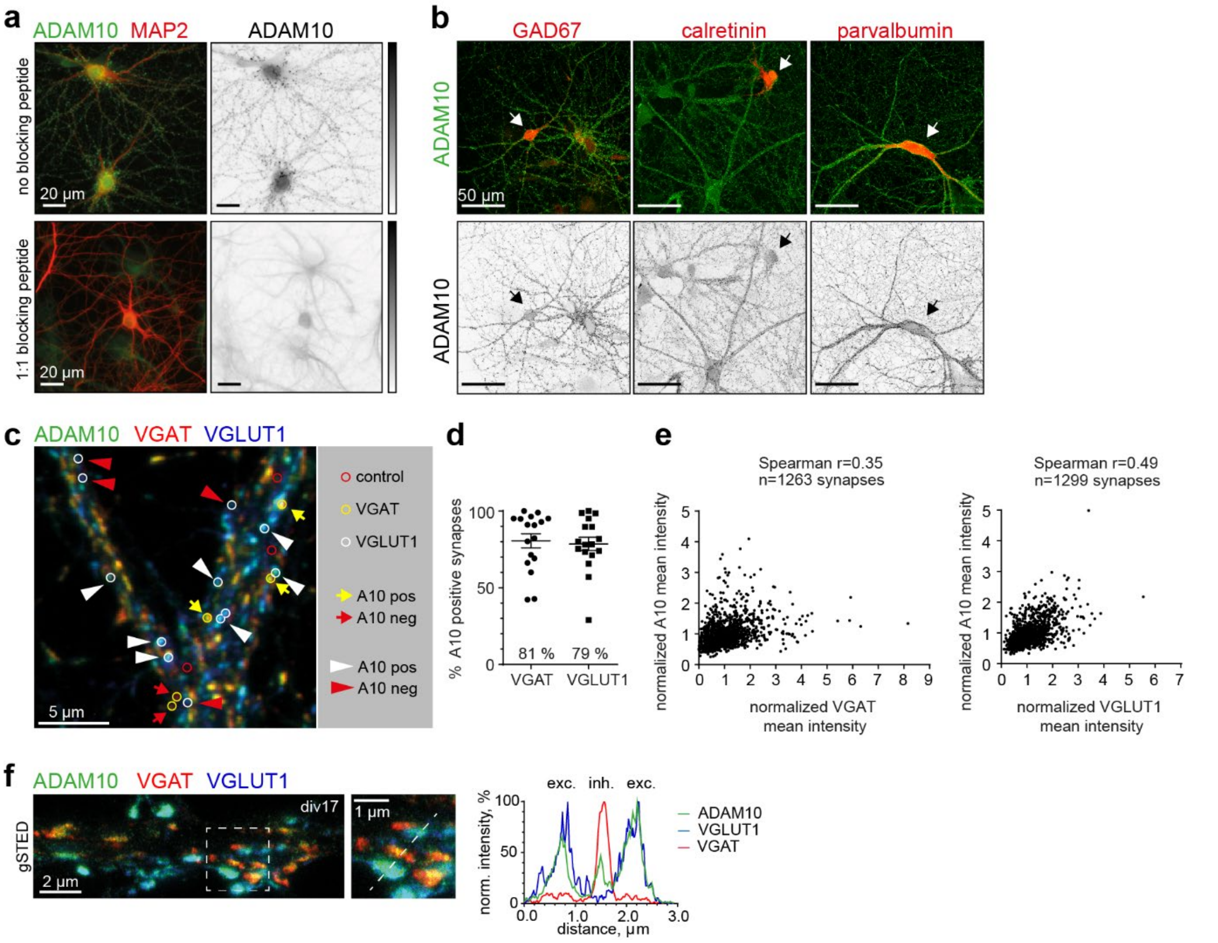


**Figure S1** Characterization of ADAM10 expression in different types of synapse and neurons

**a)** Additional anti-ADAM10-antibody characterization: Primary hippocampal rat neurons at div21 were fixed and stained for ADAM10. Note the strong signal reduction upon preincubation of the antibody with a blocking peptide (1:1 ratio).

**b)** ADAM10 is expressed in different types of interneurons. Representative maximum projections of confocal images of primary rat hippocampal cultures (div17) stained for the indicated interneuronal markers (red) and ADAM10 (green). Arrows indicate ADAM10-positive interneurons.

**c)** Representative average projections of confocal images of div17 hippocampal neuron stained with antibodies against the synaptic markers VGAT (inhibitory, red), VGLUT1 (excitatory, blue) and ADAM10 (green). A10: ADAM10.

**d)** Quantification of ADAM10-positive inhibitory and excitatory synapses labelled by VGAT and VGLUT1, respectively. Synapses are defined as ADAM10-positive if the ROIs contain more than 2 times the ADAM10 intensity than the average control ROI within the dendrite. n=17 images from 2 coverslips of 2 independent cultures. Data are represented as mean +/- SEM.

**e)** ADAM10 intensities correlate with inhibitory (VGAT) and excitatory (VGLUT1) marker intensities. n=1263 inhibitory and n=1299 excitatory synapses in 17 images from 2 coverslips of 2 independent cultures.

**f)** Representative gated STED image of hippocampal primary neurons at div17 stained for ADAM10 (green), VGLUT1 (blue) and VGAT (red). Line scans (right) show ADAM10 colocalizing with both, VGLUT1 and VGAT. A 0.5 px Gaussian blur filter was applied to the gSTED image to remove speckle noise without major changes in signal intensities. Right: Line scans of indicated synapses. exc: excitatory, inh: inhibitory synapse.

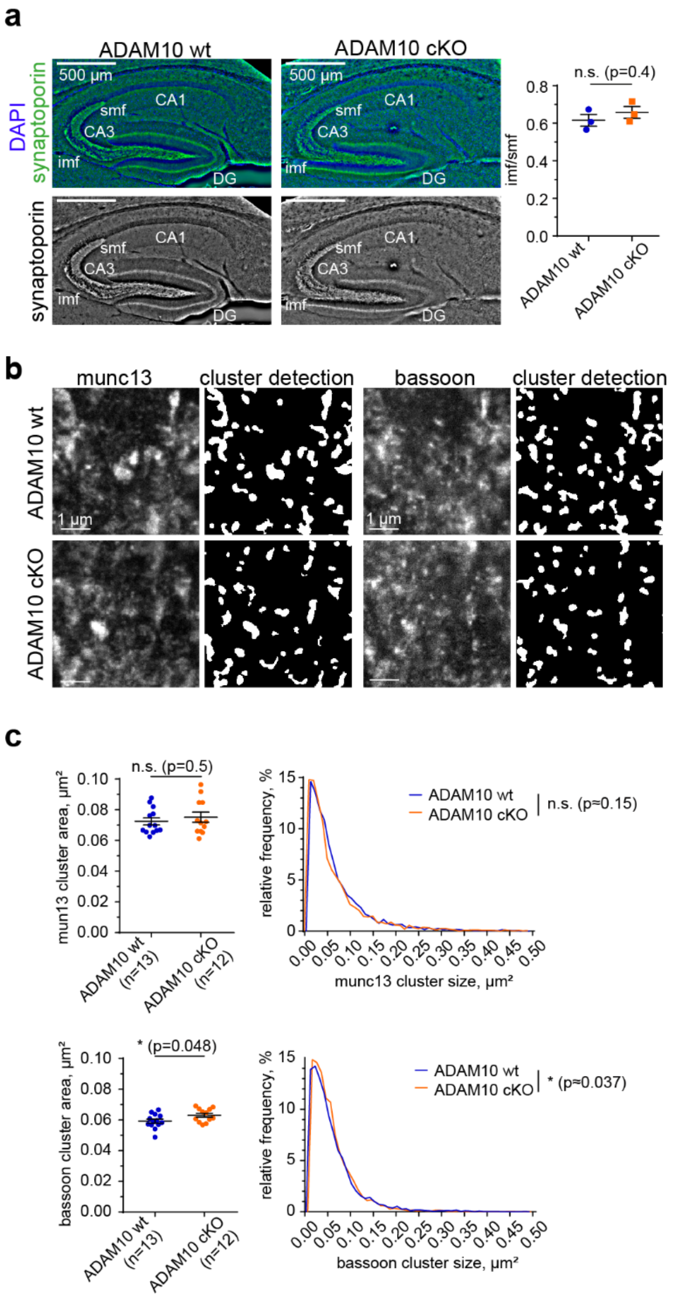


**Figure S2** Characterization of ADAM10 cKO mossy fiber boutons

**a)** Gross morphology of DG-CA3 mossy fiber tracts is unchanged in P18 A10 cKO mice. Example immunohistochemistry of sagittal cryo-sections stained for the mossy fiber marker synaptoporin and quantification of suprapyramidal mossy fiber (smf) / infrapyramidal mossy fiber (imf) length ratio. n=3 animals (3 sections each). 2-tailed unpaired Student's t-test. p=0.4. Data are represented as mean +/- SEM.

**b)** Representative STED image of mossy fiber boutons of ADAM10 wt and cKO animals (P19-21) stained for munc13 and bassoon, with corresponding detected clusters.

**c)** Quantification of average munc13 and bassoon clusters in mossy fiber boutons of ADAM10 wt and cKO. ADAM10 cKO mice show no change in munc13 presynaptic cluster area and a small increase in bassoon cluster size. n= 13 (wt) and n=12 (cKO) analyzed images of 2 sections of 2 animals each. 2-tailed unpaired Student‘s t-test. Data are represented as mean +/- SEM. Individual cluster size distributions. munc13: n= 3765 (wt), n= 3548 clusters, $p \approx 0.15$- bassoon: n= 4826 (wt), n= 4348 clusters, $p \approx 0.0372$ Kolmogorov-Smirnov test.

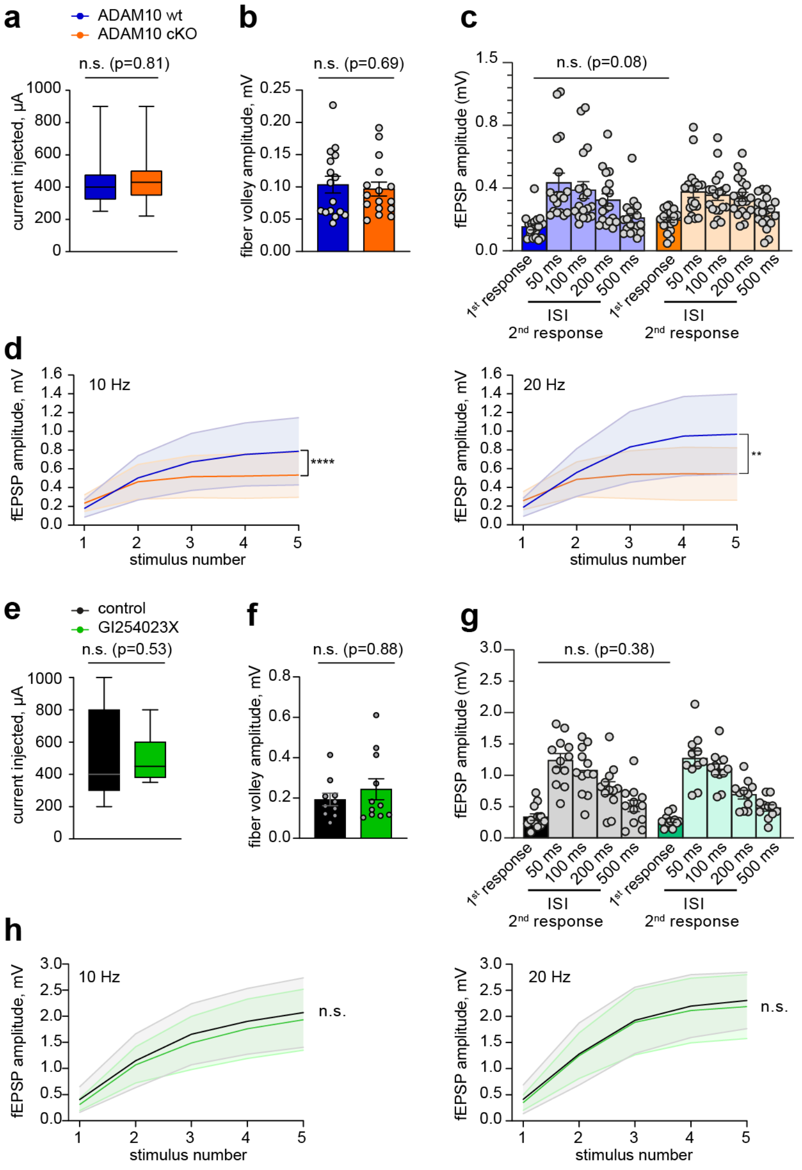


**Figure S3** Additional characterization of ADAM10-dependent mossy fiber facilitation

**a-d)** Mossy fiber plasticity of ADAM10 wt and cKO animals.

**a)** Box plot showing the electrical current required to generate action potentials in wt and cKO slices. There is no difference between genotypes. 2-tailed Student's unpaired t-test. p= 0.8115. n=13 slices (wt); n=16 slices (ADAM10 cKO) from 5 mice each. Data represented as median, 25$^{th}$ and 75$^{th}$ percentiles, minimum and maximum value.

**b)** Amplitude of fiber volleys indicating the activation of presynaptic fibers. There is no difference between genotypes. 2-tailed Student's unpaired t-test. p=0.6917. n=16 slices from 5 mice each. Data are presented as mean +/- SEM.

**c)** Amplitude of field excitatory postsynaptic potentials (fEPSP) after the first stimulus was delivered. ADAM10 cKO mice show unchanged initial neurotransmitter release. 2-tailed Student's unpaired t-test. *p=0.0766. n=18 slices (wt); n=19 slices (ADAM10 cKO) from 5 mice each. Data are presented as mean +/- SEM.

**d)** Plot of fEPSP amplitude upon train facilitation at 10 Hz and 20 Hz. ADAM10 cKO slices show an impaired response to train stimulation. 2-way repeated measures ANOVA: p<0.0001 (interaction effect, 10 Hz), p=0.0038 (20 Hz). n=18 slices; n=19 slices (ADAM10 cKO) from 5 mice each. Data are represented as mean +/- SD.

**e-h)** Mossy fiber plasticity of wt animals with or without ADAM10 inhibitor GI254023X treatment.

**e)** Box plot showing the intensity currents required to generate action potentials in wt slices with or without ADAM10 inhibitor treatment. 2-tailed Student's unpaired t-test (p=0.5279). n = 11 slices from 3 mice each. Data represented as median, $25^{th}$ and $75^{th}$ percentiles, minimum and maximum value.

**f)** Amplitudes of fiber volleys indicating the activation of presynaptic fibers. There is no effect of ADAM10 inhibitor treatment. 2-tailed Student's unpaired t-test (p=0.4188). n = 10 slices (control); n = 11 slices (GI254023X) from 3 mice each. Data are represented as mean +/-SEM.

**g)** Amplitude of the first and the second fEPSP responses at different frequencies of wt slices with or without ADAM10 inhibitor treatment. Application of GI254023X ADAM10 inhibitor has no effect on initial synaptic response. 2-tailed unpaired Student's t-test (p=0.38). n=12 slices (control); n=11 slices (GI254023X) from 3 mice each. Data are represented as mean +/-SEM.

**h)** Plot of fEPSP amplitude upon train facilitation at 10 Hz and 20 Hz. Slices treated with ADAM10 inhibitor do not show changes in response to train stimulation. 2-way repeated measures ANOVA p=0.5349 (10 Hz), p=0.7456 (20 Hz). n=12 slices (control), n=11 slices (GI254023X) from 3 mice each. Data are represented as mean +/- SD.

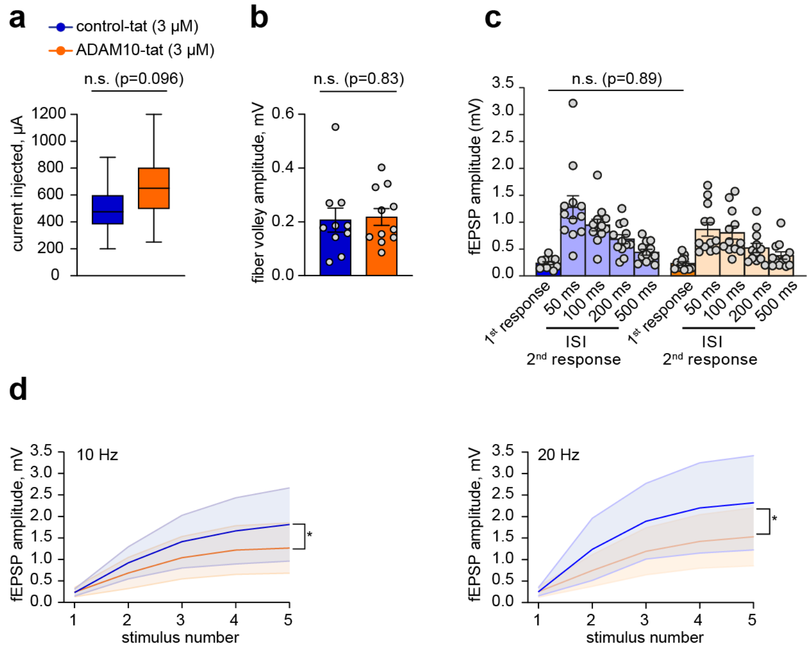


**Figure S4** Additional characterization of mossy fiber facilitation upon ADAM10 C-terminal interference

**a)** Box plot showing the electrical current applied to generate action potentials. The current injected is not significantly different between the two treatments. 2-tailed unpaired Student's t-test (p=0.1461). n=12 slices from 3 mice each. Data represented as median, 25$^{th}$ and 75$^{th}$ percentiles, minimum and maximum value.

**b)** Amplitude of fiber volleys indicating the activation of presynaptic fibers. No difference between treatments. 2-tailed unpaired Student's t-test. p=0.834. n=11 slices (ctr-tat), n=10 slices (ADAM10-tat) from 3 mice each. Data are presented as mean +/- SEM.

**c)** Amplitude of the first and the second fEPSP responses at different frequencies in wt slices treated with tat-peptides. ADAM10-tat-peptide treatment does not change the initial neurotransmitter release. 2-tailed Student's unpaired t-test. p=0.89. n=12 slices from 3 mice each. Data are presented as mean +/- SEM.

**d)** Plot of average fEPSP amplitude upon train facilitation at 10 Hz and 20 Hz. Treatment with ADAM10-tat-peptide leads to a decrease in response to train stimulation. 2-way repeated measures ANOVA. * p=0.0143 (10 Hz, interaction effect), p=0.0391 (20 Hz). n=12 slices from 3 mice each. Data are represented as mean +/-SD.
